# Structural basis for VIPP1 oligomerization and maintenance of thylakoid membrane integrity

**DOI:** 10.1101/2020.08.11.243204

**Authors:** Tilak Kumar Gupta, Sven Klumpe, Karin Gries, Steffen Heinz, Wojciech Wietrzynski, Norikazu Ohnishi, Justus Niemeyer, Miroslava Schaffer, Anna Rast, Mike Strauss, Jürgen M. Plitzko, Wolfgang Baumeister, Till Rudack, Wataru Sakamoto, Jörg Nickelsen, Jan M. Schuller, Michael Schroda, Benjamin D. Engel

## Abstract

Vesicle-inducing protein in plastids (VIPP1) is essential for the biogenesis and maintenance of thylakoid membranes, which transform light into life. However, it is unknown how VIPP1 performs its vital membrane-shaping function. Here, we use cryo-electron microscopy to determine structures of cyanobacterial VIPP1 rings, revealing how VIPP1 monomers flex and interweave to form basket-like assemblies of different symmetries. Three VIPP1 monomers together coordinate a non-canonical nucleotide binding pocket that is required for VIPP1 oligomerization. Inside the ring’s lumen, amphipathic helices from each monomer align to form large hydrophobic columns, enabling VIPP1 to bind and curve membranes. *In vivo* point mutations in these hydrophobic surfaces cause extreme thylakoid swelling under high light, indicating an essential role of VIPP1 lipid binding in resisting stress-induced damage. Our study provides a structural basis for understanding how the oligomerization of VIPP1 drives the biogenesis of thylakoid membranes and protects these life-giving membranes from environmental stress.

## Introduction

Thylakoid membranes orchestrate the light-dependent reactions of oxygenic photosynthesis, splitting water to generate the Earth’s oxygen and using the electrons and H^+^ liberated from this reaction to produce NADPH and ATP. These bioenergetic molecules are then used to remove CO_2_ from the atmosphere and convert it into the sugar that feeds Earth’s heterotrophic life. Thus, most of the energy flowing through our planet’s complex web of life originates in the intricate thylakoid membranes of plants, algae, and cyanobacteria. As climate change begins to exert increasing stress on these organisms, thylakoid architecture must be maintained to ensure the foundation of life on our planet.

Despite the central importance of thylakoids to our biosphere, we know very little about how these sheet-like membranes are constructed or how their integrity is maintained under environmental stress. Vesicle-inducing protein in plastids (VIPP1, also known as IM30) co-evolved with thylakoids and is believed to play a key role in thylakoid biogenesis and homeostasis (Li et al., 1994; Vothknecht et al., 2011). Complete knockout of VIPP1 blocks thylakoid assembly and is lethal (Zhang et al., 2012). Reduction of VIPP1 expression in the cyanobacterium *Synechocystis* sp. PCC6803 (Westphal et al., 2001; Fuhrmann et al., 2009b), the cyanobacterium *Synechococcus* sp. PCC7002 (Zhang et al., 2014), the eukaryotic green alga *Chlamydomonas reinhardtii* (Nordhues et al., 2012), and the vascular plant *Arabidopsis thaliana* (Kroll et al., 2001; Aseeva et al., 2007) strongly impairs the biogenesis of thylakoid membranes, resulting in lower levels of the membrane-embedded photosystems and a corresponding drop in photosynthetic activity. VIPP1 knockdown was also observed to cause dramatic thylakoid swelling in *Arabidopsis* and *Chlamydomonas* exposed to high light (Nordhues et al., 2012; Zhang et al., 2012), indicating that VIPP1 helps thylakoids cope with acute stress. In support of this function, overexpression of VIPP1 improved the recovery of *Arabidopsis* from heat stress (Zhang et al., 2016).

*In vitro*, VIPP1 self-assembles into large homo-oligomers of variable symmetry: *Synechocystis* VIPP1 (*syn*VIPP1) and *Arabidopsis* VIPP1 (*at*VIPP1) primarily form rings (Aseeva et al., 2004; Fuhrmann et al., 2009a; Zhang et al., 2016; Saur et al., 2017), whereas *Chlamydomonas* VIPP1 (*cr*VIPP1) makes long helical rods (Liu et al., 2007; Theis et al., 2019). Higher-order VIPP1 oligomers likely also form *in vivo*. In *Arabidopsis*, VIPP1-GFP accumulates in puncta and larger filament-like structures (Aseeva et al., 2004; Zhang et al., 2012; Zhang et al., 2016). When *Synechocystis* and *Synechococcus* are grown in high light, VIPP1-GFP forms stable puncta that have restricted mobility, likely due to membrane interactions (Bryan et al., 2014). Under moderate light, VIPP1-GFP is more dynamic and readily exchanges between soluble and punctate pools (Gutu et al., 2018). VIPP1 has no predicted transmembrane domains, yet it has a strong affinity for lipids *in vitro* and *in vivo* that is mediated by an N-terminal amphiphilic helix (Li et al., 1994; Kroll et al., 2001; Otters et al., 2013; McDonald et al., 2017). *In vitro*, *syn*VIPP1 rings have been shown to bind liposomes and induce fusion (Hennig et al., 2015), while *cr*VIPP1 rods encapsulate liposomes (Theis et al., 2019). Adding to the enigmatic complexity of VIPP1 function, *at*VIPP1 and *syn*VIPP1 were recently reported to have GTPase activity, despite the lack of a canonical GTP-binding domain (Ohnishi et al., 2018; Junglas et al., 2020).

## Results

### High-resolution model of VIPP1 oligomerization

A mechanistic understanding of the many described functions of VIPP1 requires high-resolution structural information. To date, the only available 3D structures of VIPP1 are negative-stain electron microscopy (EM) densities that lack the resolution to discern individual VIPP1 monomers (i.e., protomers) (Saur et al., 2017). Therefore, we determined structures of heterologously expressed *syn*VIPP1 rings by single particle cryo-electron microscopy (cryo-EM). Iterative classification and refinement of ~337,000 particles from 8,120 images yielded five different cryo-EM density maps corresponding to VIPP1 rings with C14, C15, C16, C17, and C18 symmetries (Figs. 1H, S1, S2) at resolutions ranging from 3.8 Å to 5.0 Å (Fig. S3). Using a combination of homology modeling, *ab initio* structure prediction, and refinement by molecular dynamics flexible fitting (MDFF), we built atomic models of the two <4 Å structures (C15 and C16) and then extended the model building to cover all five symmetries (Fig. S4A-B).

**Figure 1.**
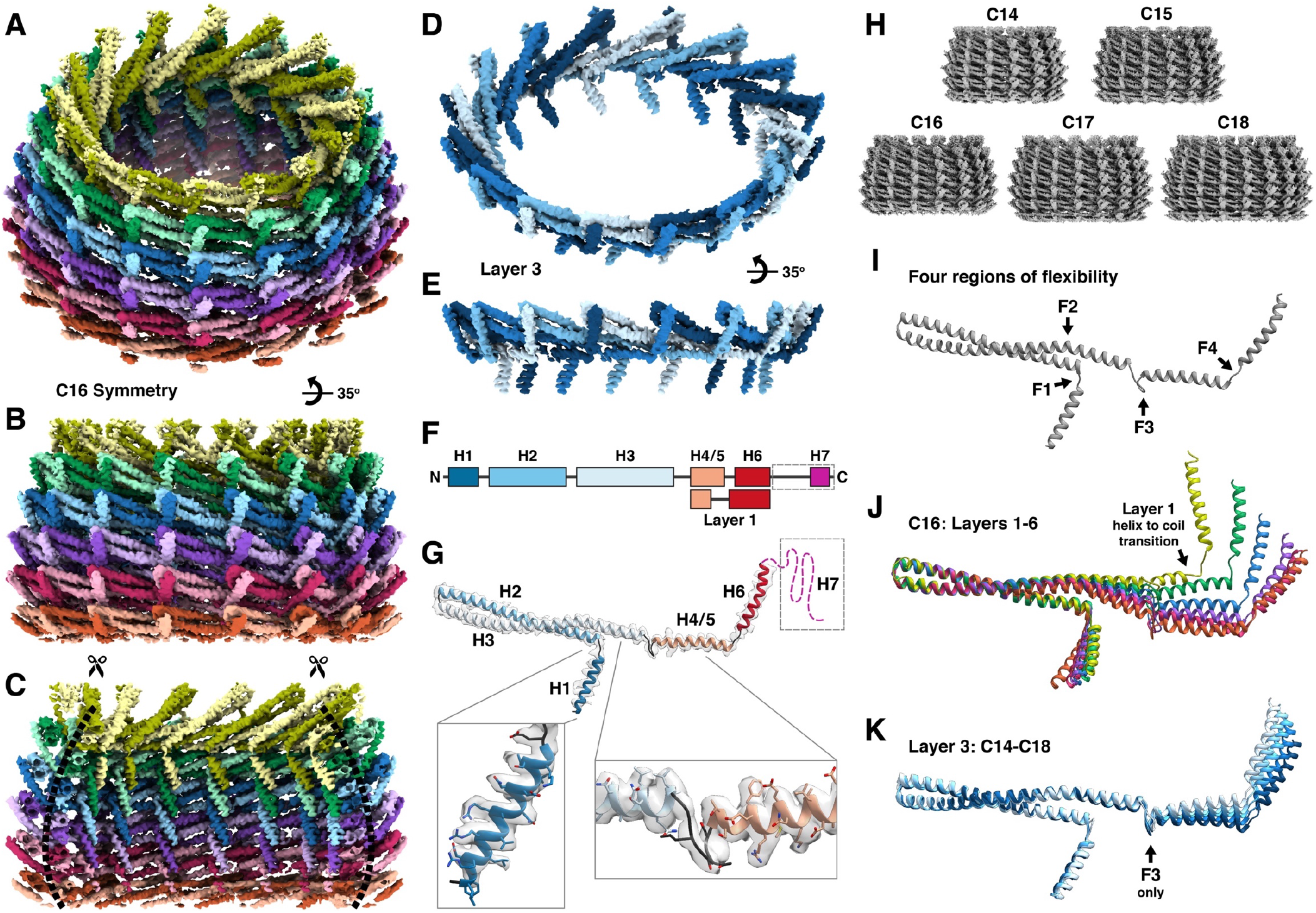
VIPP1 monomers interweave and flex to form basket-like ring structures. **(A)** Inclined and **(B)** side views of the cryo-EM density map for a *syn*VIPP1 ring with C16 symmetry. The ring consists of six layers shown in different colors (layer 1: olive green, layer 6: orange). Two shades of each color highlight the interwoven structure. **(C)** Cut-open view of the ring reveals how the H1 helices of each monomer align to form vertical columns that face toward the lumen. **(D)** Inclined and **(E)** side views of layer 3, shown in isolation to visualize the way VIPP1 monomers (in four shades of blue) extend and interact. **(F)**Schematic diagram of the VIPP1 monomer’s secondary structure elements (helices H1-H7 in colored rectangles) and **(G)** the corresponding regions on the VIPP1 molecular model, fit into the EM density (transparent white). The C-terminal domain (dashed line box), which includes H7, was not resolved in the EM density map, likely due to high flexibility. Magnified view of two regions of the monomer structure (H1 and the loop connecting H3 with H4), showing the fit of the VIPP1 model into the EM density map. **(H)** Cryo-EM VIPP1 ring structures for five symmetries (C14-C18). **(I)** Comparing the molecular models of these structures (see Figs. S4, S7) revealed four regions of flexibility in the VIPP1 monomer (F1-F4). **(J)** Superposition of the VIPP1 monomers from each layer of the C16 ring. F1-F4 all flex to assemble the ring. The N-terminal side of H4/5 is remodeled into a coil in layer 1 (see Fig. S8). **(K)** Superposition of the VIPP1 monomers from layer 3 of each symmetry (C14: darkest blue, C18: lightest blues). Only the F3 loop flexes to accommodate different ring symmetries.

The oligomeric structure of the *syn*VIPP1 ring resembles a tightly interwoven basket (Fig. 1A-C, Movie 1). Contrary to a previous model that positions the VIPP1 monomers vertically, parallel to the ring’s central axis (Saur et al., 2017), the monomers are arranged horizontally into stacked layers (Fig. 1D-E). Six layers were resolved in C14-C16, whereas C17 and C18 contained seven layers. The rings were tapered on both ends, with a smaller diameter at the top (layer 1) than the bottom (layer 6 or 7) (Fig. S4F). The VIPP1 protein is predicted to contain seven helices (H1-H7; Fig. 1F) (Otters et al., 2013; Zhang et al., 2016). Our density maps resolve five helices, with the predicted H4 and H5 domains forming a single helix, which we have named H4/5 to maintain consistent nomenclature with previous studies (Fig. 1G). The N-terminal H1 domains of each layer align with each other to form vertical columns that face the lumen of the ring (Fig. 1C). H2 and H3 make a coiled-coil hairpin, which is connected to H4/5 by a loop of seven amino acids (Fig. 1G). H6 protrudes on the outer surface of the ring (Fig. 1B). Within a layer, H3 and H4/5 together extend to pass behind two neighboring monomers, then H6 points upwards and binds the H2/H3 hairpin of the third neighboring monomer, thereby forming the interwoven structure (Fig. 1D-E). In addition, the luminal H1 columns stabilize the ring from the inside and help hold the stacked layers together. In total, each VIPP1 monomer can interact with up to 16 other monomers from three layers (Figs. S5, S6).

H7 was unresolved in our density maps, presumably due to flexibility. This small C-terminal helix with a predicted amphipathic structure is connected to H6 by an intrinsically disordered linker of ~25 amino acids. Based on the position of H6 on the outer wall of the ring, it is clear that H7 extends outside the ring. Thus, one can envision the complete VIPP1 ring as a “hairy basket” decorated with many flexible H7 domains. This structure is consistent with the dispensability of H7 for VIPP1 oligomerization (Liu et al., 2007; Otters et al., 2013; Hennig et al., 2017) and ideally positions H7 to interact with external factors in the cellular environment, such as membranes or chaperones, which may regulate the VIPP1 dynamics (Zhang et al., 2016). H1-H6 are homologous to bacterial phage shock protein A (PspA) (Westphal et al., 2001; Otters et al., 2013), which also forms an oligomeric ring (Hankamer et al., 2004), and the heterologous expression of VIPP1 can functionally complement the deletion of PspA in *Escherichia coli* (DeLisa et al., 2004; Zhang et al., 2012). However, PspA cannot compensate for VIPP1 deficiency (Zhang et al., 2016), underscoring the thylakoid-specific role of H7.

Comparing our five *syn*VIPP1 structures (Figs. 1H, S4A-B) revealed four regions of flexibility in the VIPP1 monomer (F1-F4) that allow it to build tapered rings of different symmetries (Fig. 1I). All four regions bend and flex to enable monomers to assemble into the different layers of a ring (Figs. 1J, S7A, S8, Movie 2). In contrast, for the same layer across the symmetries (e.g., layer 3 in C14-C18), the flexibility primarily comes from region F3 (Figs. 1K, S7B, Movie 3). This loop apparently enables *syn*VIPP1 rings to expand to reach higher symmetries with increased diameter.

### VIPP1 rings assemble a novel nucleotide-binding pocket

To check if unmodeled density was present, we calculated difference maps between the cryo-EM densities and the VIPP1 molecular models (Figs. 2, 3). We observed a small difference density between layers 1 and 2 (Fig. 2A-B) that could accommodate the structure of a bound nucleotide (Fig. 2C). Comparison to nucleotide densities from other cryo-EM structures (Banerjee et al., 2016) showed a better match to ADP than the larger density of ATP*γ*S (Fig. S9A). To confirm the presence of nucleotide in the *syn*VIPP1 structure, we subjected the purified protein to reversed-phase ion-pair high-performance liquid chromatography (RPIP-HPLC) coupled to electrospray ionization mass spectroscopy (ESI-MS). We detected a molecular mass corresponding to ADP (Fig. S10A-B), which we reasoned might come from the ATP wash performed during protein purification. When we sequentially washed with ATP followed by GTP, we detected ADP and GDP in the purified VIPP1 rings (Fig. 2F), indicating that VIPP1 may be able to bind and hydrolyze both ATP and GTP.

**Figure 2.**
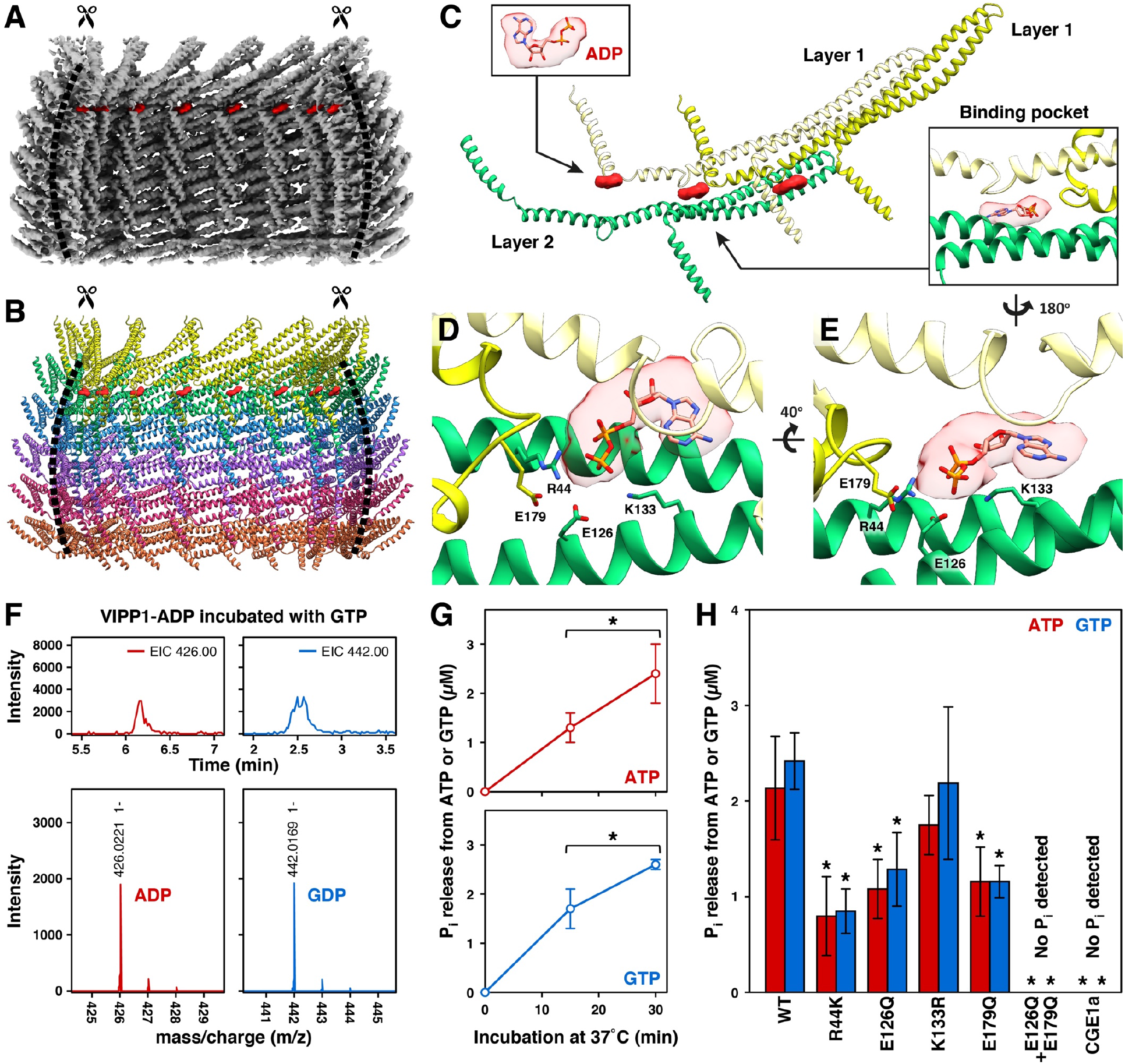
Nucleotide binding sites are formed at the interface between three VIPP1 monomers. **(A-B)** Cut-open side views of the **(A)** C16 cryo-EM density map and **(B)** molecular model, with extra EM densities in red. **(C)** The extra densities and three VIPP1 monomers extracted from the model of the complete ring (layer 1: yellow, layer 2: green). Insets show fitting of the ADP structure into the extra density (upper left) and a nucleotide binding pocket formed at the interface between three monomers (lower right). **(D-E)** Close-up views of the nucleotide binding pocket from the **(D)** top and **(E)** side. Side chains and labels are displayed for residues predicted by the model to position the nucleotide (R44, K133) and coordinate nucleophilic attack on the *γ*-phosphate. **(F)** RPIP-HPLC profiles (top panels) and subsequent ESI-MS (bottom panels) of wild-type *syn*VIPP1 following incubation with GTP. Some of the bound ADP (see Fig. S10A-B) was exchanged for GDP. **(G)** *In vitro* nucleotide hydrolysis assays showing that wild-type *syn*VIPP1 causes phosphate release from both ATP and GTP. **(H)** *In vitro* ATP and GTP hydrolysis by wild-type *syn*VIPP1 compared to point mutations in the nucleotide binding pocket (double mutant is indicated with a “+”). CGE1a was used as a negative control. This nucleotide exchange factor of chloroplast Hsp70 does not possess ATPase activity and was purified using the same protocol as VIPP1. Error bars are standard deviation from 6-10 replicates, asterisks indicate a significant change (p<0.05, Welch’s t-test).

**Figure 3.**
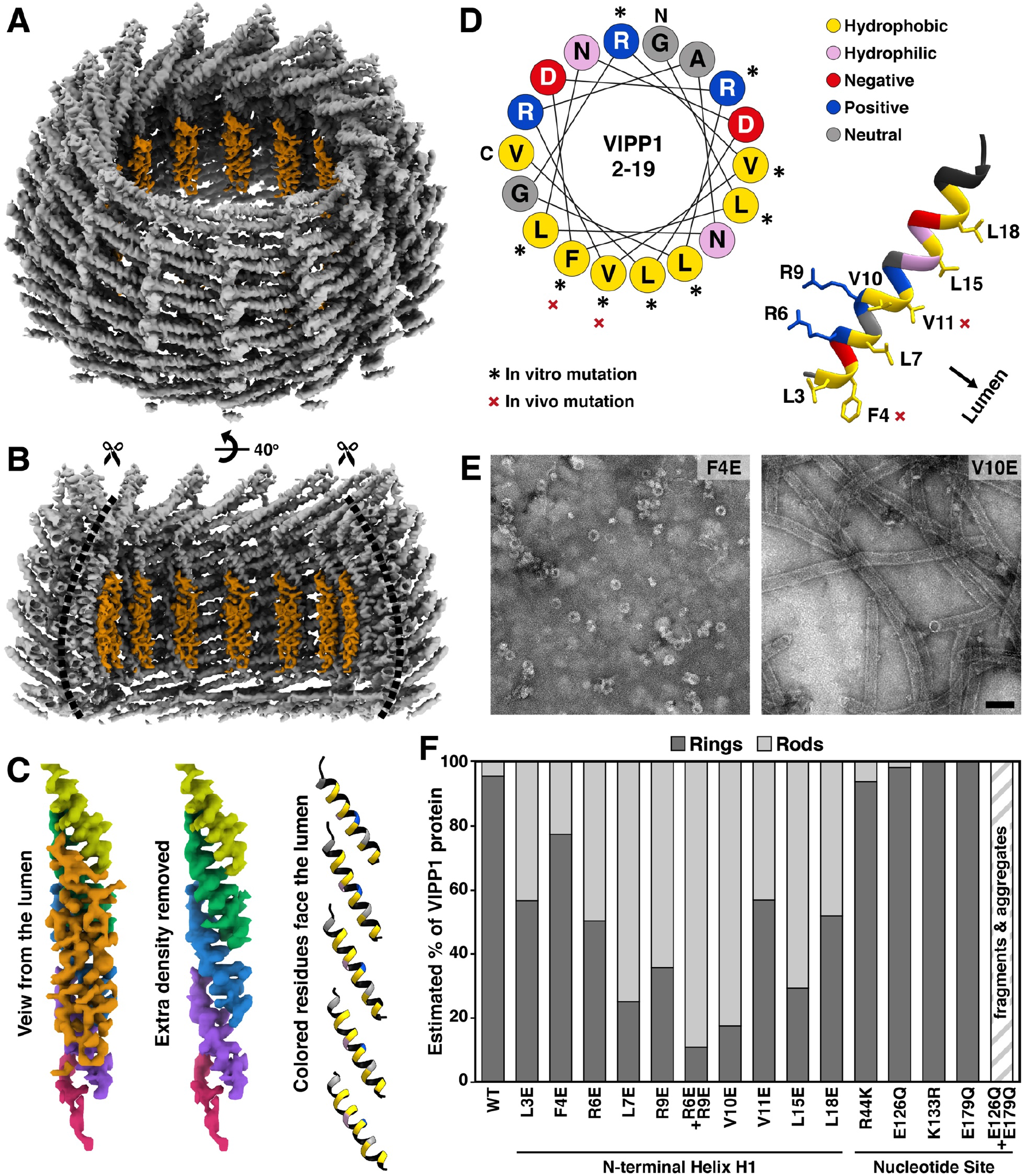
Columns of H1 helices form large hydrophobic surfaces on the luminal face of the VIPP1 ring. **(A)** Inclined and **(B)** side views of the C15 cryo-EM density map, with extra EM densities shown in orange. **(C)** Luminal view of a column of N-terminal H1 helices, shown as EM density (colored by layer according to Fig. 1, left column with orange extra density, middle column with extra density removed) and as a molecular model (right column, only the lumen-facing residues are colored according to panel D). Yellow hydrophobic residues face the ring lumen. Helix H1 of layer 5 (magenta) is poorly resolved due to flexible interactions with the unanchored, and thus unresolved, H1 of layer 6. **(D)** Helical wheel plot of residues 2-19 (left) and corresponding molecular model (right), showing the amphipathic structure of H1. *In vitro* mutations (see Figs. S10E, S11) are indicated by an asterisk and shown with labeled side chains on the molecular model. *In vivo* mutations (see Figs. 4–6, S13–S14) are indicated by a red “x”. **(E)** Example negative stain EM images for two *in vitro* mutant proteins and **(F)** the estimated percentage of protein assembled into rings or rods (see Figs. S11, S12) for all *in vitro* mutants in this study, including nucleotide pocket mutants from Fig. 2. Scale bar: 100 nm.

The VIPP1 nucleotide binding site is formed by bringing together different regions of three VIPP1 monomers, two from layer 1 and one from layer 2 (Fig. 2C). To our knowledge, this tripartite oligomeric mechanism of nucleotide pocket formation is novel. The nucleotide is accommodated by flexibility in region F4 of layer 1, with the C-terminal portion of H4/5 opening into a short coil (Figs. 1J, S8). We fit an ADP molecule into the EM density and refined the molecular model by interactive MDFF (Goh et al., 2016), revealing characteristic structural and charge distribution features of a nucleotide binding pocket (Figs. 2D-E, S9B). In our refined model, Lys133 and Arg44 from layer 2 use their positive charges to clamp the phosphates of the nucleotide. Glu126 from layer 2 as well as Glu179 on the opened coil of layer 1 (Fig. S8) are well positioned to orient a water molecule for nucleophilic attack on the *γ*-phosphate. When the nucleotide densities from the C15, C16, and C17 rings were viewed at a lower threshold, each displayed a small additional density that could correspond to Mg^2+^ (Fig. S9A), a standard feature of NTPase pockets that helps coordinate the nucleotide. VIPP1 lacks one common element of ATP and GTP binding sites, the Walker A (ATPases) or P-Loop (GTPases) motif (Walker et al., 1982). However, there is no universal fingerprint for nucleotide binding, and other known NTPases also lack this motif (Saraste et al., 1990).

We next performed *in vitro* nucleotide hydrolysis assays and observed that purified *syn*VIPP1 and *cr*VIPP1 can hydrolyze both ATP and GTP (Figs. 2G, S10C-D), consistent with previous reports of GTPase activity in *syn*VIPP1 and *at*VIPP1 (Ohnishi et al., 2018; Junglas et al., 2020). Point mutations in *syn*VIPP1 Arg44, Glu126, and Glu179 reduced the phosphate release for both ATP and GTP (Fig. 2H), supporting our structural model of the nucleotide binding pocket. Double mutation of Glu126 and Glu179 abolished phosphate release, further corroborating that these residues coordinate hydrolysis. Mutation of Lys133 to Arg did not have a significant effect. However, residue 133 is an Arg in *cr*VIPP1 and PspA, possibly explaining the tolerance of the *syn*VIPP1 nucleotide pocket to K133R substitution. Whereas all the *syn*VIPP1 proteins with single point mutations in the nucleotide pocket produced ring structures, the G126E+G179E double mutant did not assemble into rings or rods, instead forming fragments and disorganized aggregates (Fig. S11). Therefore, we conclude that nucleotide hydrolysis is required for *syn*VIPP1 oligomerization into ring structures.

### Structural basis for VIPP1 lipid binding

A second difference density not accounted for by VIPP1 monomers was found inside the lumen of the ring specifically along the H1 columns (Fig. 3A-C). Previous studies have determined that H1 is required for lipid binding (Otters et al., 2013; McDonald et al., 2015; McDonald et al., 2017), and our structure clearly reveals the mechanism of this interaction. H1 is an amphipathic helix that orients its hydrophilic side toward the VIPP1 ring wall and its hydrophobic side towards the lumen (Fig. 3D). As a result, the H1 columns form large hydrophobic surfaces on the inside of the VIPP1 ring (Fig. 3C). These hydrophobic H1 columns explain how liposomes are drawn into *cr*VIPP1 rods (Theis et al., 2019) and how *syn*VIPP1 rings can mediate liposome fusion (Hennig et al., 2015). The extra density seen along the H1 columns is likely endogenous lipid, which is commonly co-purified with VIPP1 from *E. coli* cells (Otters et al., 2013).

We made structure-directed point mutations to the H1 hydrophobic interface, as well as to Arg residues proposed to impart specificity to negatively-charged lipids (McDonald et al., 2015; McDonald et al., 2017) (Fig. 3D). To our surprise, all of these H1 mutations caused *syn*VIPP1 to form more rod structures (Figs. 3E-F, S11, S12). Thus, binding of H1 to lipids may influence how VIPP1 oligomeric assemblies form, tuning the equilibrium between rings and rods. Previous work has shown that H1 is unstructured in solution and acquires α-helical structure upon interaction with lipids (McDonald et al., 2017). Given the stabilizing interactions between H1 columns and the ring wall (Figs. 1C, S4C, S5C), it is possible that lipid binding induces a structural change in H1, which then shifts the equilibrium between tapered ring and parallel rod architecture. Interestingly, several H1 mutations also showed reduced phosphate release from ATP and GTP (Fig. S10E), indicating a potential allosteric effect between lipid binding and nucleotide hydrolysis.

### VIPP1 lipid interaction protects thylakoids under high-light stress

Based on our high-resolution structure, we made *in vivo* point mutations in two conserved H1 residues previously shown to mediate lipid binding in PspA (F4E and V11E) (Jovanovic et al., 2014; McDonald et al., 2017) and then examined the effects on thylakoid architecture inside native cells using *in situ* cryo-electron tomography (Asano et al., 2016; Rast et al., 2019). In low light (30 μmol photons m^−2^s^−1^), both mutants grew slightly slower than a wild-type control strain (WT) (Figs. 4A, S13A-B). In high light (200 μmol photons m^−2^s^−1^), the growth rate of F4E slightly decreased and V11E failed to grow at all. To test the acute effects of high-light stress, liquid cell cultures were grown to log-phase in low light and then transferred to high light for 24 hours. VIPP1 levels were reduced in both mutant strains compared to WT (Figs. 4B, S13C-F), indicating that VIPP1 may be destabilized *in vivo* when lipid binding is impaired. However, upon shifting to high light, the mutant and WT strains all showed a similar ~2-fold increase in VIPP1 protein levels (Fig. 4B). In WT cells, thylakoid architecture remained relatively unchanged in high light, with thylakoid lumen width expanding mildly from ~6.5 to ~9 nm (Fig. 4C-E and L, Movie 4). Both mutant strains showed a moderate degree of thylakoid swelling in low light, but when shifted to high light, they both experienced extreme swelling (~12 to ~34 nm in F4E, ~16 to ~47 nm in V11E) (Fig. 4K-L, Movies 5-6). The severity of thylakoid swelling varied between individual cells (Figs. 4F and I, S14). As thylakoids swell, high curvature is maintained at thylakoid tips, presumably by the CurT protein (Heinz et al., 2016). Under prolonged stress, the tip curvature is lost, resulting in the formation of large round vesicles covered in membrane-bound ribosomes and phycobilisomes, markers of their thylakoid origin. While both mutants experience extreme thylakoid swelling (Fig. 4L), it is interesting that F4E survives high-light exposure better than V11E (Fig. 4A), despite lower VIPP1 protein levels (Fig. 4B). This result underscores the importance of Val11, which is centrally positioned in the H1 hydrophobic surface (Fig. 3D). It remains to be discovered how high-light stress causes thylakoid swelling and how the interaction of VIPP1 with membranes counteracts this architectural change.

**Figure 4.**
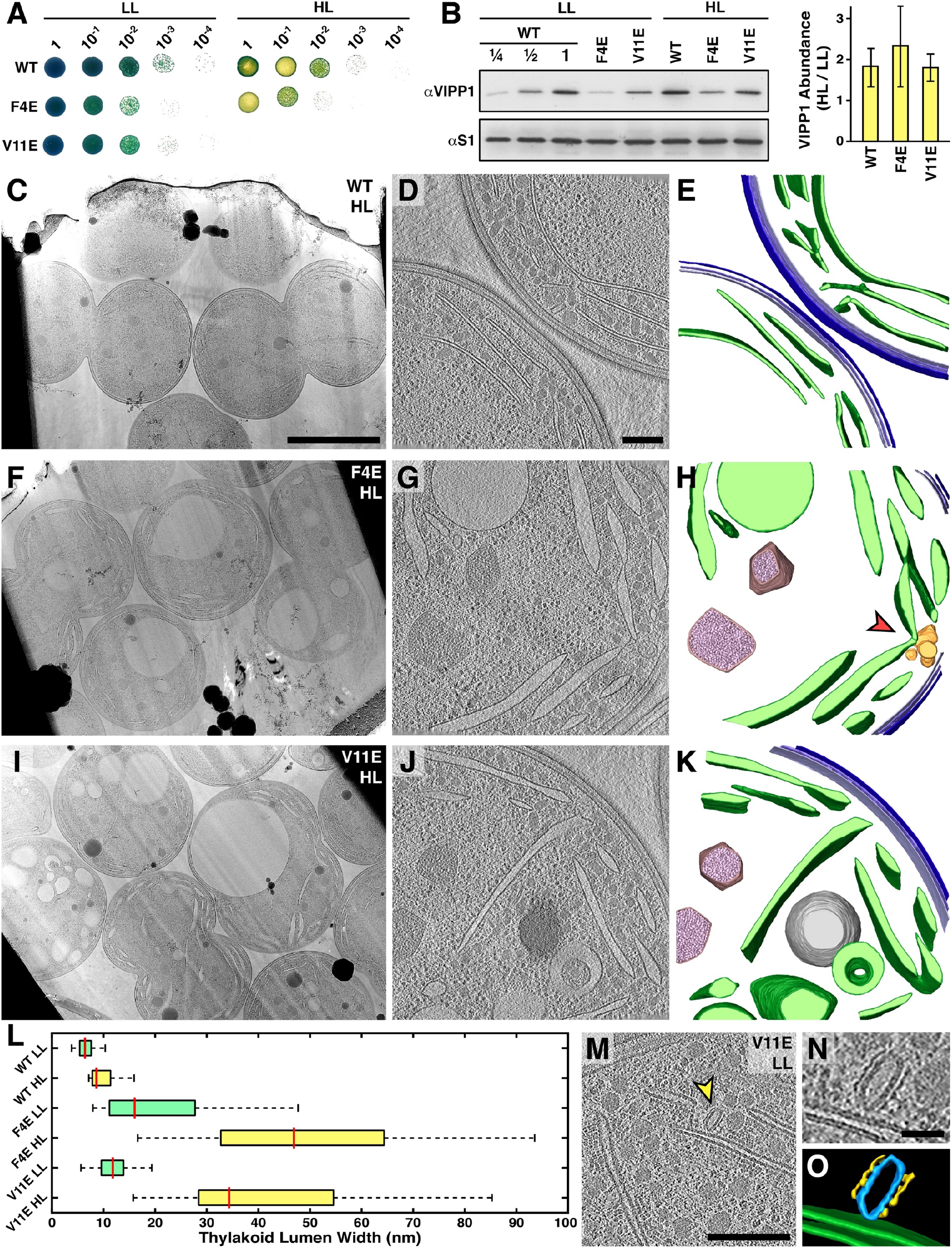
VIPP1 lipid interaction maintains thylakoid architecture under high-light stress. **(A)** Spot growth tests of WT (control where a kanamycin resistance cassette was introduced upstream of the vipp1 gene at the same position as for the mutants), F4E, and V11E stains after five days on agar plates in low light (LL, 30 μmol photons m^−2^s^−1^) and high light (HL, 200 μmol photons m^−2^s^−1^). **(B)** Western blot analysis of VIPP1 protein levels in liquid cultures at LL and after switch to HL for 24 hours. The S1 ribosomal protein was used as a loading control. Relative change between the two light conditions is quantified in the plot on the right, showing mean and standard deviation (error bars) from three independent experiments. Additional data in Fig. S13. **(C-K)** *In situ* cryo-electron tomography of WT (C-E), F4E (F-H), and V11E (I-K) cells in liquid culture after switch to HL for 24 hours. Left panels show multiple cells in low magnification overviews, central panels show 2D slices through tomographic volumes (additional examples in Fig. S14), right panels show 3D segmentations of the tomograms (dark green: thylakoid membranes; light green: thylakoid lumen; three shades of blue: outer membrane, peptidoglycan layer, and plasma membrane; brown/pink: carboxysomes; grey: lipid droplet; yellow: defective convergence membrane). Red arrowhead indicates abnormal convergence zone architecture (more examples in Fig. 5). **(L)** For each strain and condition, quantification of median thylakoid lumen width (red line), along with 25%-75% percentiles (box) and 1.5x interquartile range (whiskers). N= 283 thylakoid membrane regions from 48 tomograms. **(M-O)** Tomographic overview, close-up, and 3D segmentation of an *in situ* structure that resembles a protein coat (yellow) encapsulating a membrane (blue), in close contact with a thylakoid (green). For more examples and comparison to *in vitro* VIPP1, see Fig. 6. Scale bars: 2 μm in C, F, I; 200 nm in D, G, J, M; 40 nm in N.

In addition to an impaired acute stress response, both mutants exhibited aberrant membrane structures at the thylakoid convergence zones, regions that have been implicated in biogenesis (Stengel et al., 2012; Rast et al., 2015). Instead of the high-curvature convergence membranes that contact the plasma membrane (Figs. 4D-E) (Rast et al., 2019), we observed clusters of round membranes that appeared to contain cytosolic material (Figs. 4G-H, 5). Similar aberrant membranes have been observed at the thylakoid convergence zones of *Chlamydomonas* VIPP1 knockdown cells (Nordhues et al., 2012), indicating an evolutionarily conserved function. Careful inspection of the tomograms also revealed structures in close perpendicular contact with thylakoid membranes that resembled membranes encapsulated by a protein coat (Figs. 4M-O, 6). While further investigation is required to confirm the molecular identity of these coated membranes, we note that structural features such as the protein coat diameter and membrane-protein distance are consistent with VIPP1 rings and rods (Fig. 6I-J). Thus, these *in situ* structures might correspond to the membrane-associated VIPP1 puncta previously observed by fluorescence microscopy (Bryan et al., 2014; Gutu et al., 2018). If these *in situ* structures are indeed VIPP1-coated membranes, their presence in the F4E and V11E strains (Figs. 4M-O, 6F-H) indicates that these point mutations do not completely ablate VIPP1 lipid binding, which is consistent with the milder phenotype compared to VIPP1 knockout.

**Figure 5.**
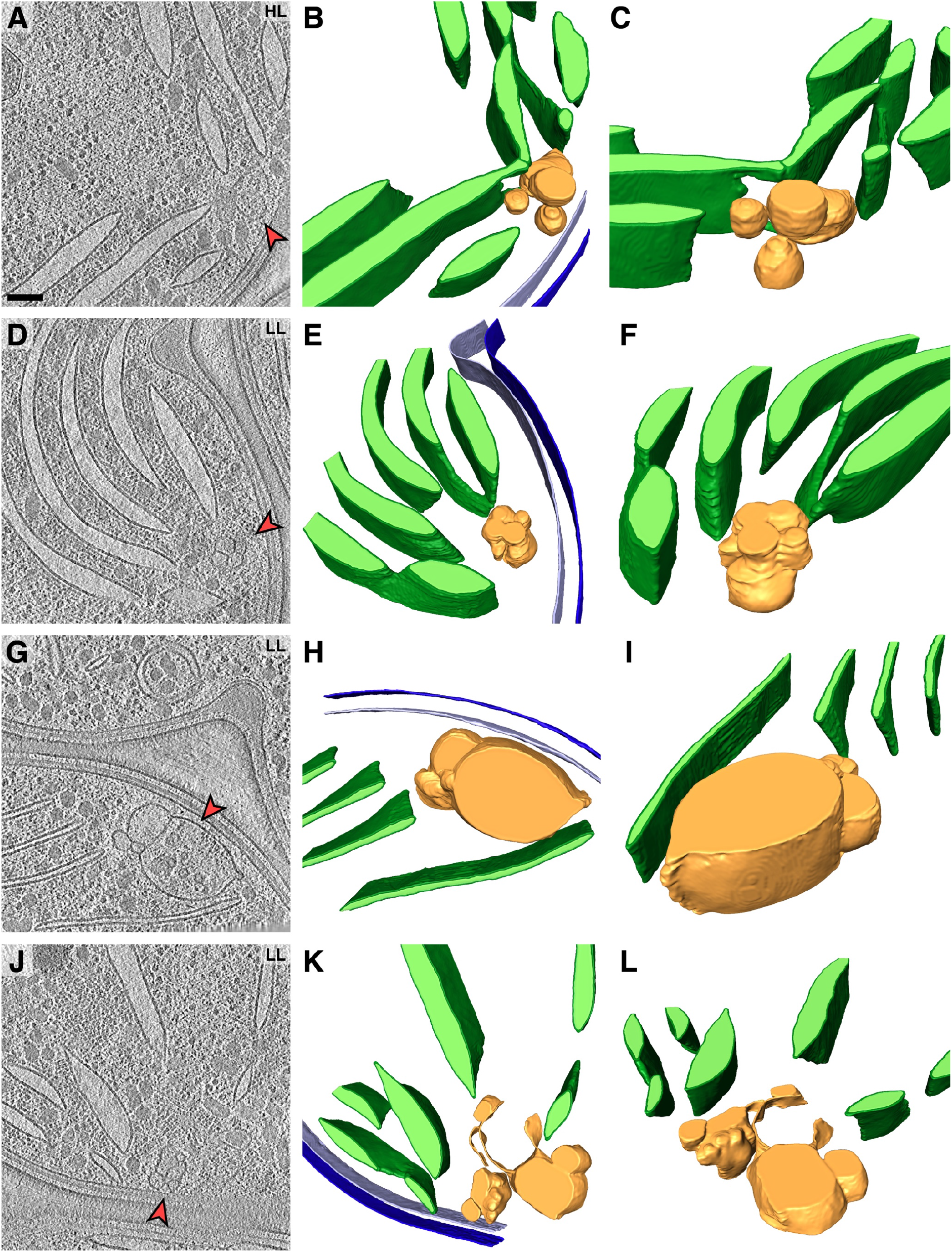
Examples of defective convergence zones in the F4E mutant. **(A,D,G,J)** 2D slices through tomographic volumes. Red arrowheads point to abnormal membranes filled with cytosolic material at convergence zones. HL: high light, LL: low light. **(B,E,H,K)** Top views and **(C,F,I,L)** inclined views showing 3D segmentations of the tomograms (dark green: thylakoid membranes; light green: thylakoid lumen; dark blue: outer membrane, light blue: plasma membrane; yellow/gold: defective convergence membranes). Panels A-C are additional views of Fig. 4G-H. For examples in the V11E mutant, see Fig. S14. Scale bar: 100 nm.

**Figure 6.**
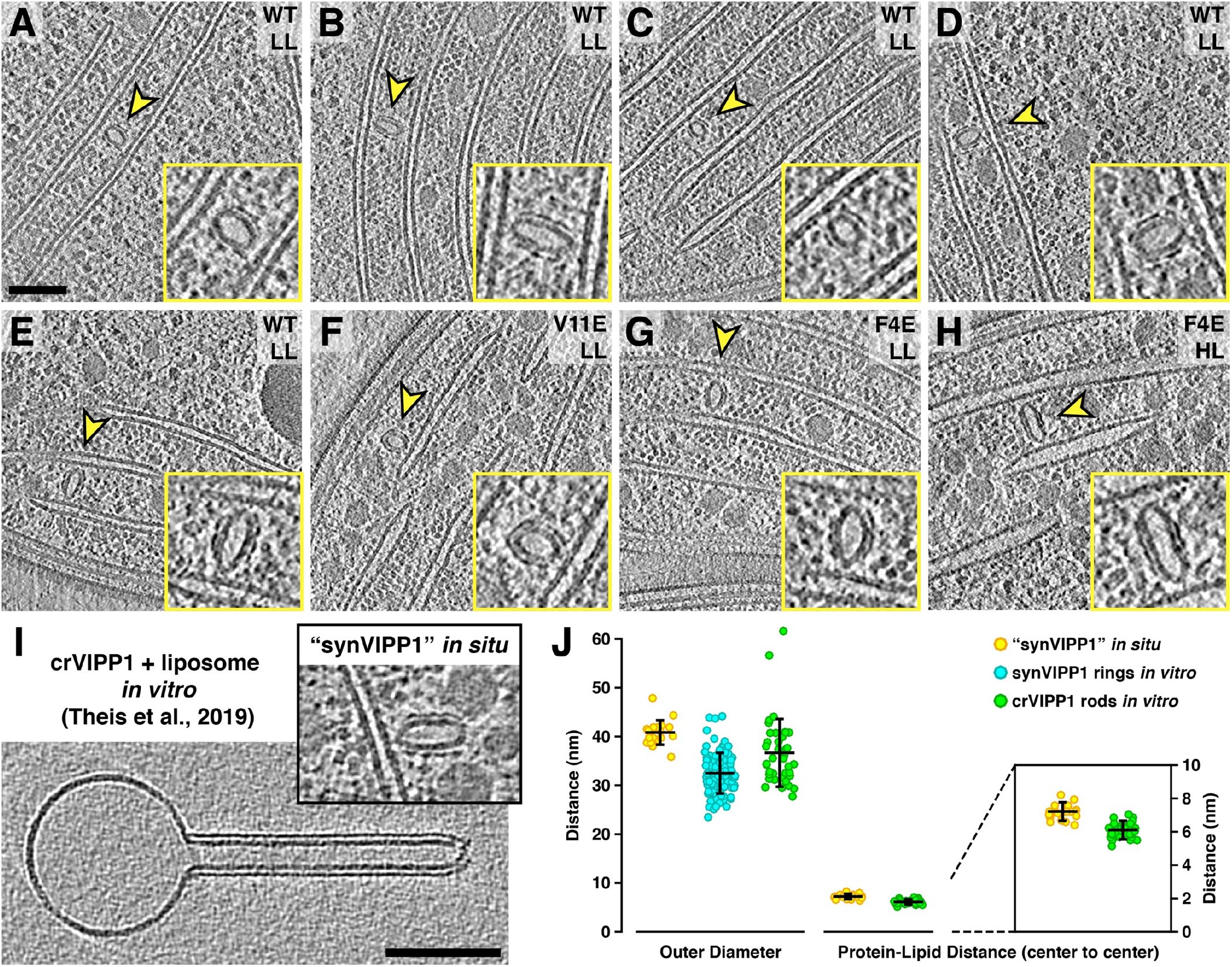
Putative *syn* VIPP1 structures visualized inside native cells by *in situ* cryo-ET and comparison of structural dimensions to *in vitro* VIPP1. **(A-H)** *In situ* tomographic slices showing membranes encapsulated in a protein coat (indicated with yellow arrowheads in the overviews and enlarged 2x in the yellow inset panels). These coated membranes were always observed closely associated with a thylakoid membrane, roughly perpendicular to the thylakoid surface. Strain and growth condition (LL: low light, HL: high light) are indicated in the top right. **(I)** Comparison of one of the *in situ* coated membrane structures (labeled “synVIPP1”) with a *cr*VIPP1 rod encapsulating a liposome *in vitro* (liposome on the left is engulfed by VIPP1 on the right; reproduced from (Theis et al., 2019)). Both images are shown at the same scale. **(J)** Measurement of the outer diameter across the short central axis of *in situ* “*syn*VIPP1” structures (yellow), *in vitro syn*VIPP1 rings (blue, from raw micrographs such as Fig. S2B), and *in vitro cr*VIPP1 rods (green) (Theis et al., 2019). For *in situ* “*syn*VIPP1” and *in vitro cr*VIPP1, the distance is also measured from the center of the protein coat density to the center of the enclosed lipid bilayer density (the inset enlarges this part of the plot). One reason for the slightly larger *in situ* distance may be that the cellular membranes are thicker than the liposomes. Horizontal black lines are mean values, error bars are standard deviations. Scale bars: 100 nm.

## Discussion

Thylakoid membranes produce the oxygen and biochemical energy that fuel most of the life on Earth. Our study describes with molecular detail how VIPP1 oligomerizes and encapsulates lipids to protect thylakoid membranes from environmental stress. Our cryo-EM structures reveal how VIPP1 monomers interweave and flex to form rings of variable symmetry. We describe a novel nucleotide pocket that is formed by bringing together different regions of three monomers. The hydrolysis of ATP or GTP in this pocket is required for VIPP1 oligomerization into rings. Furthermore, our study reveals the structural basis for VIPP1 lipid binding via its N-terminal amphipathic helix H1, which we link to the *in vivo* protection of thylakoid architecture from high-light stress. We find that point mutations in H1 shift the equilibrium of oligomerization from rings to rods, indicating that VIPP1 is on a tipping point between these two structures, which may impart flexibility to adopt varied architectures *in vivo*.

### The *in vivo* relevance of VIPP1 rings and rods

The VIPP1 rings and rods described in our study were formed from protein exogenously expressed in *E. coli*. However, several lines of evidence support the *in vivo* relevance of oligomeric VIPP1 ring and rod structures: **1)**The ring-forming helices H1-H6 of VIPP1 are so closely related to the bacterial stress-response protein PspA that VIPP1 can complement the deletion of PspA in *E. coli* (DeLisa et al., 2004; Zhang et al., 2012). Importantly, PspA isolated from its native bacterial source also forms rings (Hankamer et al., 2004), supporting the idea that rings function within the cell. **2)**The F4 and V11 residues that we selected for our *in vivo* VIPP1 mutagenesis experiments (Figs. 4–5) are conserved with PspA. Previous studies have shown that the F4E and V11E mutations each prevent PspA from binding lipids (Jovanovic et al., 2014; McDonald et al., 2017). This lipid-binding function is consistent with the positions of F4 and V11 that we see in our VIPP1 ring structure, on the H1 hydrophobic surface facing the bound material in the ring lumen (Fig. 3D). **3)** Using *in situ* cryo-ET, we show that introducing V11E and F4E mutations into endogenous VIPP1 causes defective convergence zone architecture and extreme thylakoid swelling under high-light conditions (Figs. 4, 5, S14). Thus, lipid binding inside the VIPP1 ring by the hydrophobic columns of aligned H1 helices likely functions *in vivo* to maintain the integrity of thylakoids, improving cell survival under environmental stress. **4)** In a previous study, we showed that the luminal surfaces of VIPP1 rods have high affinity for lipids and can encapsulate liposomes, sucking them up like a straw (Theis et al., 2019). Remarkably, we see structures inside the cell that closely mirror this previously described *in vitro* phenomenon (Fig. 6). The cellular structures look like rings or short rods encapsulating membranes. They have diameters that match the *in vitro cr*VIPP1 rods and *syn*VIPP1 rings. But instead of sucking up liposomes, these putative *in situ* VIPP1 structures may be sucking up thylakoids. These structures are always in close perpendicular contact with thylakoids, and the curvature of the encapsulated membrane is often pointed where it contacts the thylakoid sheet, hinting that lipids are being exchanged between the two membranes (Figs. 4M-O, 6B,E,G).

### The role of VIPP1 in thylakoid biogenesis

The molecular structures described in our study provide a starting point for unraveling the *in vivo* function of VIPP1 in thylakoid biogenesis. Thylakoid convergence zones, where thylakoids merge into high-curvature membranes that make close contact with the plasma membrane, have previously been implicated in the biogenesis of thylakoids and photosystem II (PSII) (Stengel et al., 2012; Rast et al., 2015; Heinz et al., 2016; Rast et al., 2019). In both low and high light, the F4E and V11E mutants had aberrant convergence zones that lacked thylakoid connectivity and contact sites with the plasma membrane. Instead, these regions frequently contained clusters of round membranes that were filled with a variety of cytosolic material, including ribosomes, glycogen granules, and phycobilisomes (Figs. 5, S14). Thus, the interaction of VIPP1 with membranes is clearly important for shaping wild-type convergence zone architecture. Interestingly, despite their defective convergence zones, the F4E and V11E mutants were able to assemble multi-layered thylakoid networks that contained relatively wild-type levels of photosynthetic proteins, including PSII (Fig. S13C-D). To better understand the role of VIPP1 in thylakoid biogenesis, it will be interesting to examine the F4E and V11E mutants during “greening”, the *de novo* assembly of thylakoids following nitrogen starvation (Klotz et al., 2016).

### VIPP1 protects thylakoids from stress-induced swelling

A major question posed by our study is how high light causes thylakoid swelling, and how VIPP1 counteracts this force to maintain thylakoid architecture (Figs. 4, S14). The light-induced swelling of thylakoids may be due to changes in luminal pH and the resulting increased osmotic pressure. It was long believed that lumen acidification by photosynthetic water splitting and proton pumping causes cations such as Na^+^ and K^+^ to be displaced from the thylakoid lumen, forcing the lumen to narrow due to osmotic water efflux (Murakami and Packer, 1970; Staehelin et al., 1990). However, a more recent study observed expansion of the thylakoid lumen when shifting *Arabidopsis* from dark to light (Kirchhoff et al., 2011). The authors proposed that lumen acidification leads to an influx of Cl^−^ anions through voltage-gated channels (Spetea and Schoefs, 2010), driving osmotic water influx and thylakoid swelling. Consistent with this study, we also observed thylakoid lumen expansion in response to increased light, so perhaps lumen acidification causes an influx of water instead of an efflux. How luminal ion balance drives osmotic swelling requires further investigation. The luminal concentration of K^+^ and Na^+^ cations may play an important role, as an H^+^/K^+^ antiporter (KEA3) has been localized to the thylakoid membranes of *Arabidopsis* (Kunz et al., 2014), and a related H^+^/Na^+^ antiporter (Nhas3) is present in *Synechocystis* thylakoids (Tsunekawa et al., 2009). Under high light, decreased luminal pH could potentially drive cation antiport into the thylakoid lumen, leading to osmotic water influx and thylakoid swelling. In addition to lumen acidification, high light also causes PSII to generate reactive oxygen species, which might promote thylakoid swelling by destabilizing thylakoid proteins and lipids.

How does VIPP1 counteract thylakoid swelling? One possibility is that VIPP1 oligomers actively work on stressed thylakoids to maintain their flat architecture. This idea is supported by fluorescence microscopy observations that VIPP1 puncta form at thylakoid membranes in response to high light (Bryan et al., 2014; Gutu et al., 2018). VIPP1 and PspA both have high affinity for anionic lipids (Kobayashi et al., 2007; Hennig et al., 2015; Theis et al., 2019) and can sense stored curvature elastic (SCE) stress, which bends the membrane and makes the bilayer more porous (McDonald et al., 2015). These two affinities may be related, as anionic lipids favor curved membranes due to repulsion between their headgroups (Hirama et al., 2017). Oligomeric PspA rings, but not monomers, have been shown to bind damaged bacterial membranes and prevent H^+^ leakage *in vitro* (Kobayashi et al., 2007). Through this interaction, PspA rings function *in vivo* to maintain the proton gradient across the bacterial plasma membrane under stress conditions (Kleerebezem et al., 1996). VIPP1 rings and rods may serve a similar function for thylakoids. Interestingly, VIPP1 membrane binding was recently observed to be enhanced by acidic conditions *in vitro* (Siebenaller et al., 2020), implying that VIPP1 may be able to sense H^+^ leakage in stressed thylakoid regions. However, it remains to be tested whether swollen thylakoids in the F4E and V11E mutants leak enough H^+^ to influence luminal pH. While the evidence is compelling for VIPP1 oligomerization at thylakoids with high SCE stress, it remains unclear how oligomeric VIPP1 stabilizes these membranes. Perhaps VIPP1 rings and rods extract lipids from damaged thylakoids, pulling the membrane sheet tight to physically resist outward-pushing swelling forces. The extraction of damaged lipids or specific lipid species, in particular anionic lipids that prefer curved membranes, might help make the thylakoids more rigid and flat. Removing specific lipids might also impair the function of thylakoid-embedded cation antiporters (van der Does et al., 2000), reducing cation influx and the resulting osmotic pressure. While this requires further investigation, we may indeed see evidence of such a mechanism *in situ*, with putative VIPP1 structures extracting lipids from thylakoid sheets (Fig. 6).

Another possible explanation is that VIPP1 plays a more constitutive role, maintaining thylakoid connectivity and lipid composition to make the thylakoid network more resistant to swelling. VIPP1 has been implicated in membrane fusion (Hennig et al., 2015), and we primarily observed the putative VIPP1 structures between two thylakoids (Fig. 6). Importantly, we observed these structures under all light intensities, not only in high-light stress. Therefore, VIPP1 may act constitutively to facilitate lipid exchange between neighboring thylakoids or even fuse these membranes to increase connectivity of the whole thylakoid network. The reduced thylakoid connectivity, defective convergence zone architecture, and absence of plasma membrane contact sites in the F4E and V11E mutants (Fig. 5) may result in altered thylakoid lipid composition that is more vulnerable to osmotic forces. Note, however, that thylakoid lipid composition in *Chlamydomonas* VIPP1 knockdown cells was previously observed to be indistinguishable from wild type (Nordhues et al., 2012).

### Nucleotide hydrolysis and the polarity of VIPP1 oligomerization

Like actin filaments and microtubules, VIPP1 rings (and presumably also rods) are polar structures that require nucleotide binding for assembly. However, unlike these cytoskeletal structures, VIPP1 rings do not retain nucleotides bound along their lengths. It remains to be described how nucleotide binding and hydrolysis drive VIPP1 oligomerization. The VIPP1 structures in our study hint at two distinct possible mechanisms depending on the polarity of oligomer assembly (Fig. 7). If layer 1 is the first to assemble, nucleotide binding and hydrolysis may only be required for initial nucleation of the ring structure, and then extension of the structure proceeds without nucleotide binding. If layer 1 is the last to assemble, it is likely that oligomerization proceeds through a cycle of nucleotide binding, hydrolysis, and release as layers are sequentially added. If VIPP1 oligomerization is nucleated on thylakoid membranes, this “hydrolysis and release” mechanism could potentially produce the thylakoid-associated putative VIPP1 structures we observe *in situ* (Fig. 6). VIPP1’s amphipathic H1 helix senses SCE stress and has high affinity for anionic lipids (Otters et al., 2013; Hennig et al., 2015; McDonald et al., 2015; McDonald et al., 2017). We propose that VIPP1 monomers interact with thylakoids via their H1 helix and concentrate in areas with high levels of SCE stress and/or anionic lipids, where they begin to oligomerize into membrane-anchored ring and rod structures. As VIPP1 rings oligomerize away from the membrane, sequentially adding layers by nucleotide hydrolysis and release, they suck up and encapsulate thylakoid lipids in the ring lumen (Fig. 7). These encapsulated membranes may be enriched in anionic lipids, due to the affinity of these lipids for both VIPP1 and curved membranes regions.

**Figure 7.**
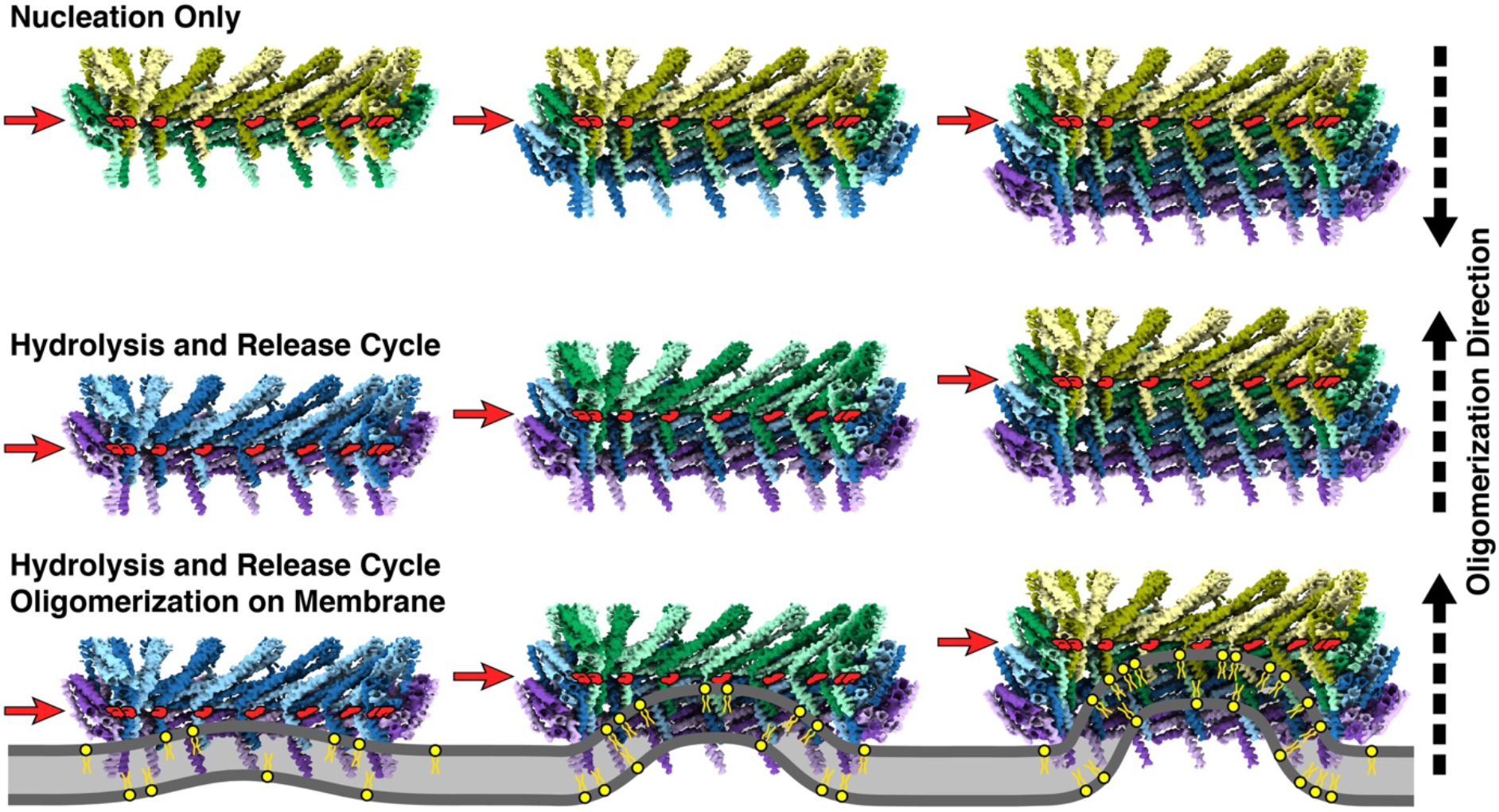
Hypothetical models for the polarized assembly of VIPP1 rings. In the “nucleation only” model (top row), ring structures are nucleated by the binding and hydrolysis of nucleotides between layers 1 and 2, the same position that ADP densities are observed in the final cryo-EM density maps (nucleotide positions are marked with red arrows). Oligomerization then proceeds sequentially to layers 3, 4, and so on, without additional nucleotide binding. In the “hydrolysis and release” model (middle row), ring structures are nucleated on the opposite end from layer 1 (nucleotide is shown between layers 3 and 4 here, but it would start between layers 5 and 6 in the C16 ring from Fig. 1). After hydrolysis, the nucleotide is released, and then more nucleotide is bound and hydrolyzed as a new layer is added (shown here between layers 2 and 3, middle panel). This cycle continues until additional layers cannot be added, presumably due to the inclination of layer 1. Regardless of the direction of oligomerization (indicated with dashed arrows on the right of the figure), the final structure looks the same in both models, with nucleotide bound between layers 1 and 2. The “hydrolysis and release” model is more consistent with oligomerization of VIPP1 on a membrane (bottom row), potentially yielding the encapsulated membrane structures seen associated with thylakoid membranes *in situ* (Fig. 6). In such a mechanism, VIPP1 monomers interact with the thylakoid membrane (grey) through their amphipathic H1 helix and begin to concentrate at regions that have high SCE stress and/or are rich in anionic lipids (yellow). VIPP1 monomers oligomerize into a membrane-anchored ring structure that extends by “hydrolysis and release”, adding new layers further and further from the membrane. As the VIPP1 ring extends, lipids are sucked from the thylakoid into the growing ring lumen. The increased membrane curvature and affinity for VIPP1 could potentially concentrate anionic lipids in this encapsulated membrane, extracting them from the thylakoid sheet.

VIPP1’s nucleotide-regulated assembly mechanism is likely conserved with PspA, which has also been observed to hydrolyze GTP (Ohnishi et al., 2018). Once assembled, VIPP1 rings and rods are rather stable but can be efficiently disassembled by HSP70B-CDJ2-CGE1 chaperones (Liu et al., 2007). The interplay of membrane interaction, nucleotide hydrolysis, and chaperone binding likely regulates the *in vivo* dynamics of VIPP1 oligomerization, as well as the balance between ring and rod structures.

### Similarities between VIPP1 and other membrane remodeling proteins

The oligomer-forming helices of VIPP1 (H1-H6) are closely conserved with PspA. Therefore, the structural mechanisms revealed in our study are almost certainly relevant to PspA’s function in maintaining plasma membrane integrity, with implications for bacterial virulence and antibiotic resistance (Darwin, 2013; Manganelli and Gennaro, 2017). In addition to PspA, VIPP1 also has some structural and functional similarities to ESCRT-III. Like VIPP1, ESCRT-III contains extended α-helices connected by flexible loops that enable it to interweave into oligomers of variable symmetry that bind and curve membranes (McCullough et al., 2015; Nguyen et al., 2020). ESCRT-III drives membrane fission in numerous cellular processes (Hurley, 2015), and like both VIPP1 and PspA, it plays important roles in membrane repair. In mammalian cells, ESCRT-III has been implicated in nuclear envelope repair (Denais et al., 2016; Raab et al., 2016) and mitotic reformation (Olmos et al., 2015; Vietri et al., 2015) as well as repair of the plasma membrane (Jimenez et al., 2014; Scheffer et al., 2014). However, VIPP1/PspA and ESCRT-III have distinct evolutionary origins, having arisen in bacteria and archaea, respectively (Lindås et al., 2008; Samson et al., 2008).

### Adaptation of thylakoid membranes to a rapidly changing climate

VIPP1 is widely conserved throughout photosynthetic organisms, from land crops that feed billions of people to marine algae and cyanobacteria that fix half of the Earth’s CO_2_. VIPP1’s essential function of maintaining thylakoid integrity under environmental stress, including excess light and heat, makes it a key player in the adaption of photosynthetic systems to our rapidly changing climate. Our study opens new opportunities for engineering plants and algae that can better resist extreme environmental conditions, helping ensure food security for the world’s growing population while fixing more atmospheric CO_2_ to combat climate change.

---

**Figure S1.**
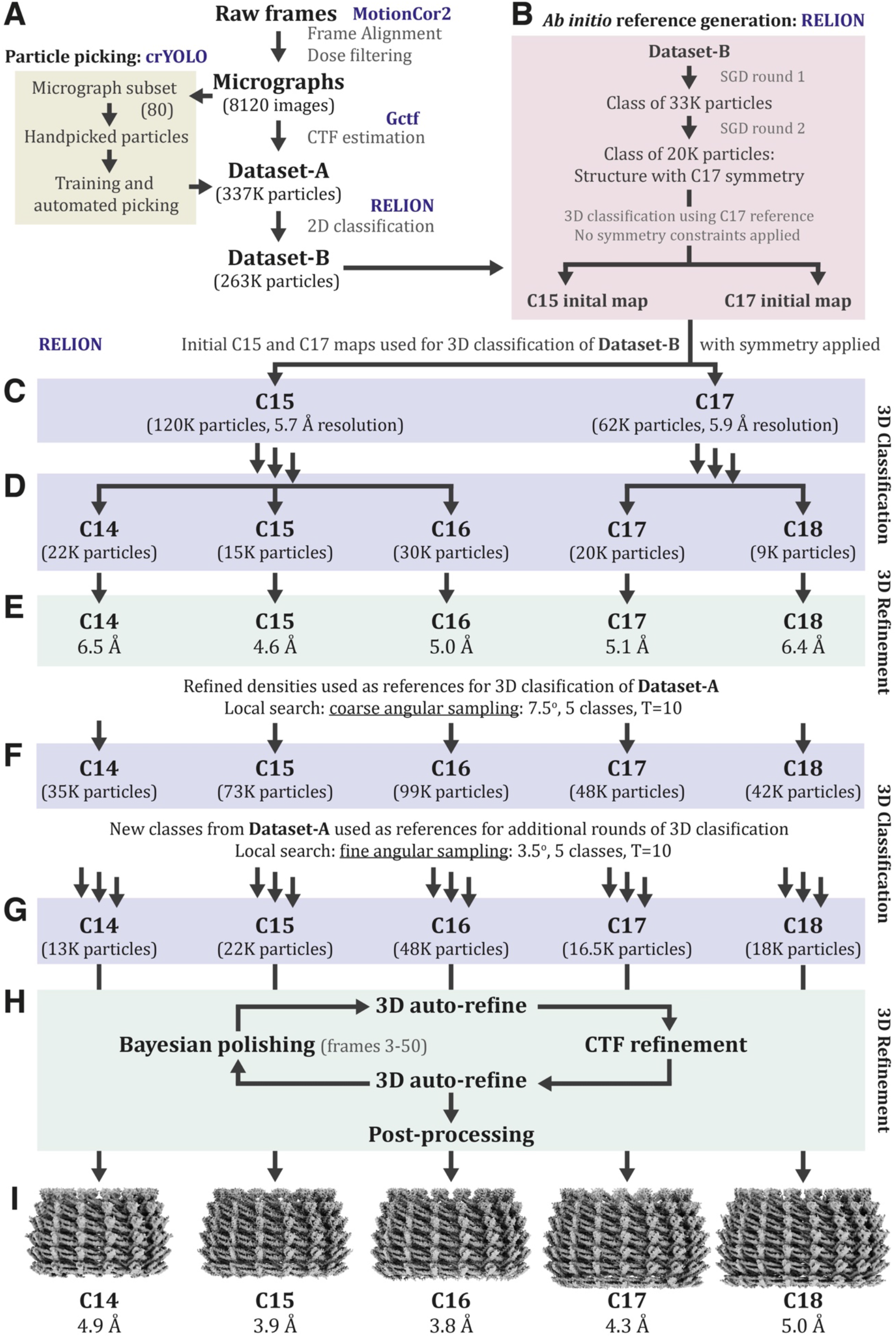
Schematic overview of the cryo-EM single particle data processing workflow. For an accompanying detailed description, please see the Methods. Blue text indicates the software used for each step. **(A)** 8120 frame stacks were aligned and dose filtered (Zheng et al., 2017), subjected to CTF estimation (Zhang, 2016), and then ~337,000 particles (Dataset-A) were extracted by deep learning-based automated picking (Wagner et al., 2019). One round of 2D classification reduced the dataset to ~263,000 particles (Dataset-B). **(B)** Stochastic gradient decent (SGD) and 3D classification were applied to a subset of Dataset-B to generate C15 and C17 starting maps *ab initio*, without prior reference (Zivanov et al., 2018). **(C)** 3D classification of the full Dataset-B using these *ab initio* references and enforced symmetry split the dataset into C15 and C17 classes. **(D)** Subsequent rounds of 3D classification yielded classes for C14-C18 symmetries. **(E)** These were refined to produce final averages for Dataset-B. **(F-G)** The Dataset-B maps of five symmetries were used for several iterative rounds of 3D classification on Dataset-A, yielding classes of C14-C18 with more particles. **(H)** 3D refinement of these classes included iterative rounds of RELION 3D auto-refinement (Scheres, 2012), per-particle defocus estimation (CTF refinement) (Zivanov et al., 2018), and per-particle motion correction and dose-weighting (Bayesian polishing) (Zivanov et al., 2019). **(I)** The resulting final EM density maps of C14-C18 rings had resolutions ranging from 3.8 Å to 5.0 Å, as judged by “gold standard” Fourier shell correlation (FSC) comparing independently refined half-maps (see Fig. S3A).

**Figure S2.**
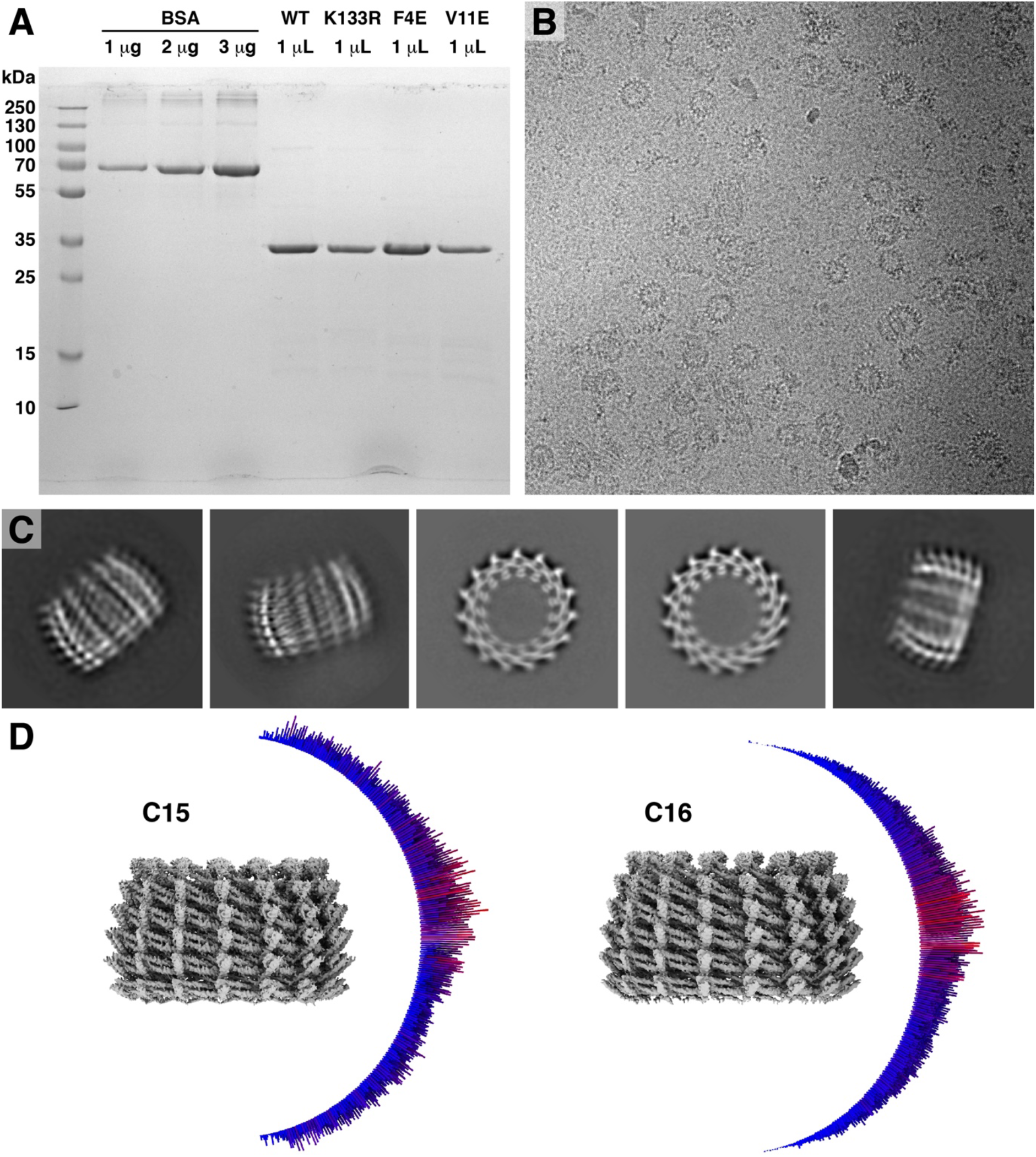
*syn* VIPP1 protein purification and steps of Cryo-EM image analysis. **(A)** Coomassie-stained SDS-PAGE of four different *syn*VIPP1 preps, run along with BSA protein concentration standards and a molecular weight ladder. VIPP1 has a molecular weight of ~29 kDa. **(B)** Example raw micrograph containing top and side views of *syn*VIPP1 particles. **(C)** Representative reference-free 2D class averages, showing side and top views. **(D)** Angular distribution of particles in the final C15 and C16 averages. More populated side views are red. Less populated oblique and top views are blue.

**Figure S3.**
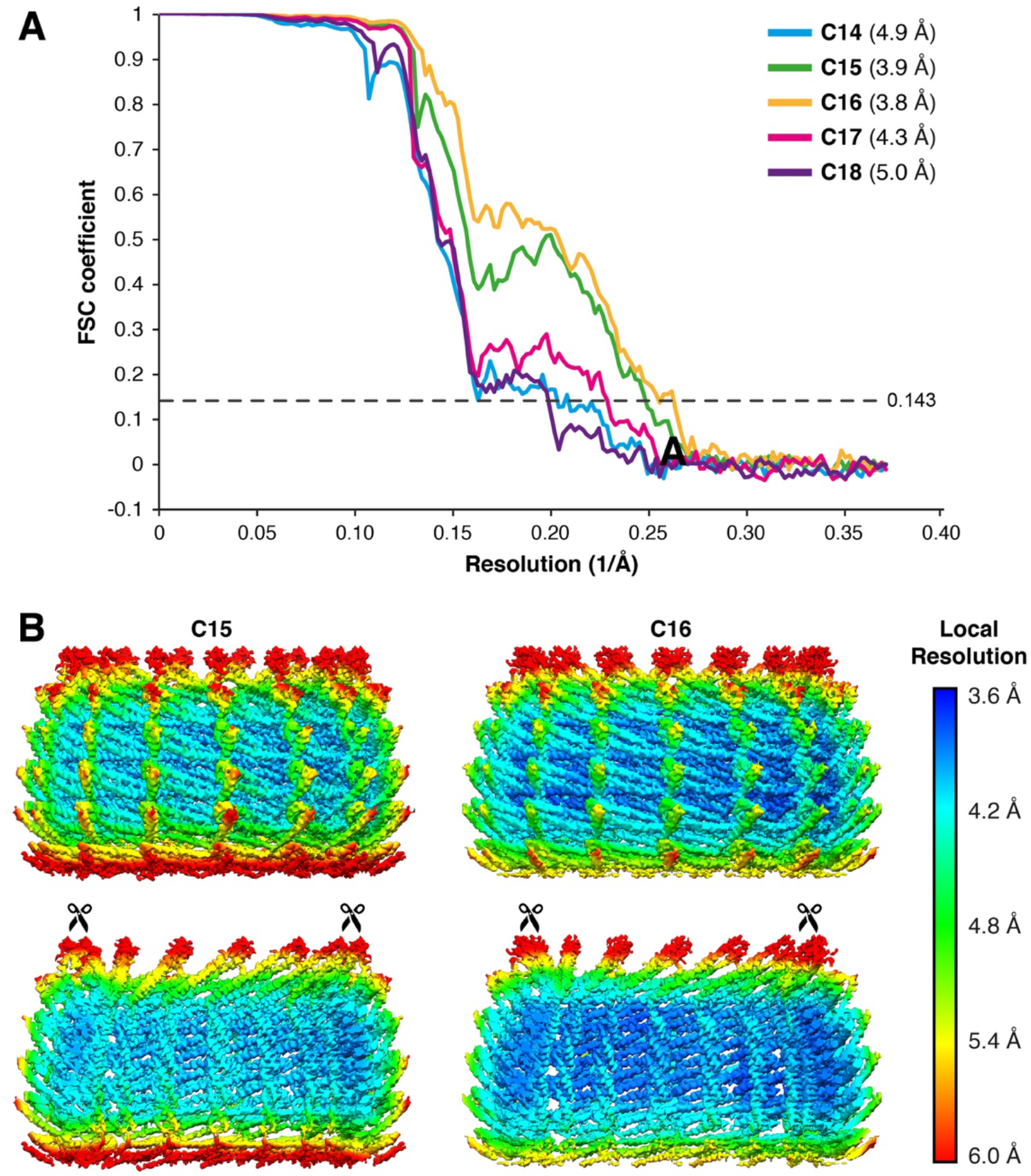
Resolution estimation of the cryo-EM structures. **(A)** “Gold-standard” Fourier shell correlation (FSC) plots for the five VIPP1 ring structures (C14-C18). Numerical resolution values were determined with the FSC=0.143 cutoff (dashed line). **(B)** Local resolution of the C15 and C16 maps, estimated in RELION with the local resolution function and plotted in UCSF Chimera. All structures are side views, with the top panels showing the outside surfaces of the rings and the bottom panels showing cut-open views of the rings that reveal the H1 helices.

**Figure S4.**
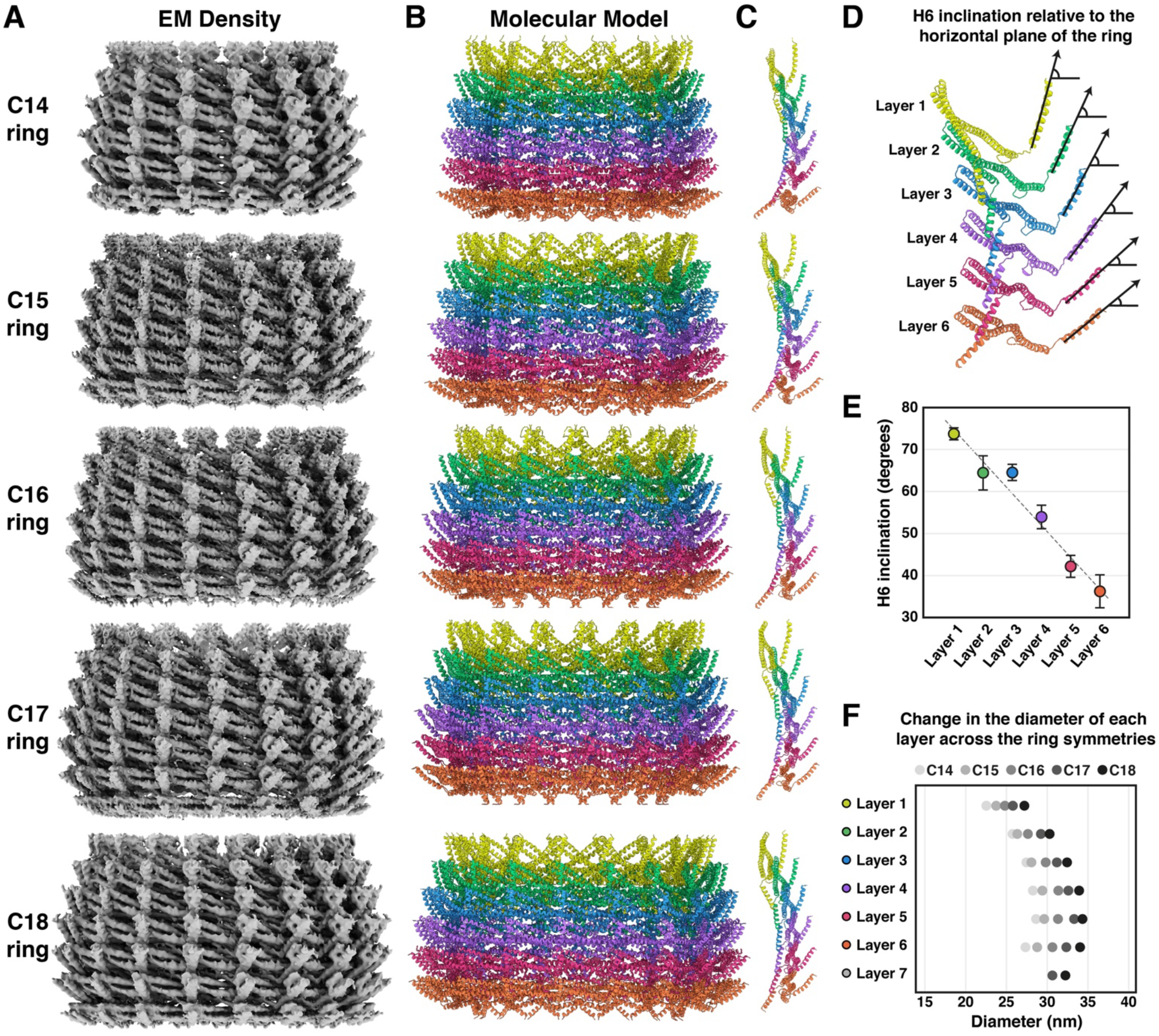
Molecular modeling for VIPP1 rings of each symmetry. **(A)** Cryo-EM density maps and **(B)** corresponding molecular models for five different symmetries of VIPP1 rings. Density maps for C14, C15 and C16 have six layers, whereas C17 and C18 have seven layers (only the first six layers from the top could be modeled due to the low resolution of layer 7). **(C)** One vertical row of monomers is displayed, showing the inclination of H1 (left side) and H6 (right side) for each ring structure. **(D-E)** For each Vipp1 structure, the inclination of H6 was measured as the angle relative to the horizontal plane of the ring. **(E)** Plot of the mean (dot) and standard deviation (error bars) from all symmetries. **(F)** Plot showing the diameter of each layer from the structures of each symmetry. Diameters were measured from the outer walls of the rings. Note that layer 7 was only observed in the C17 and C18 rings.

**Figure S5.**
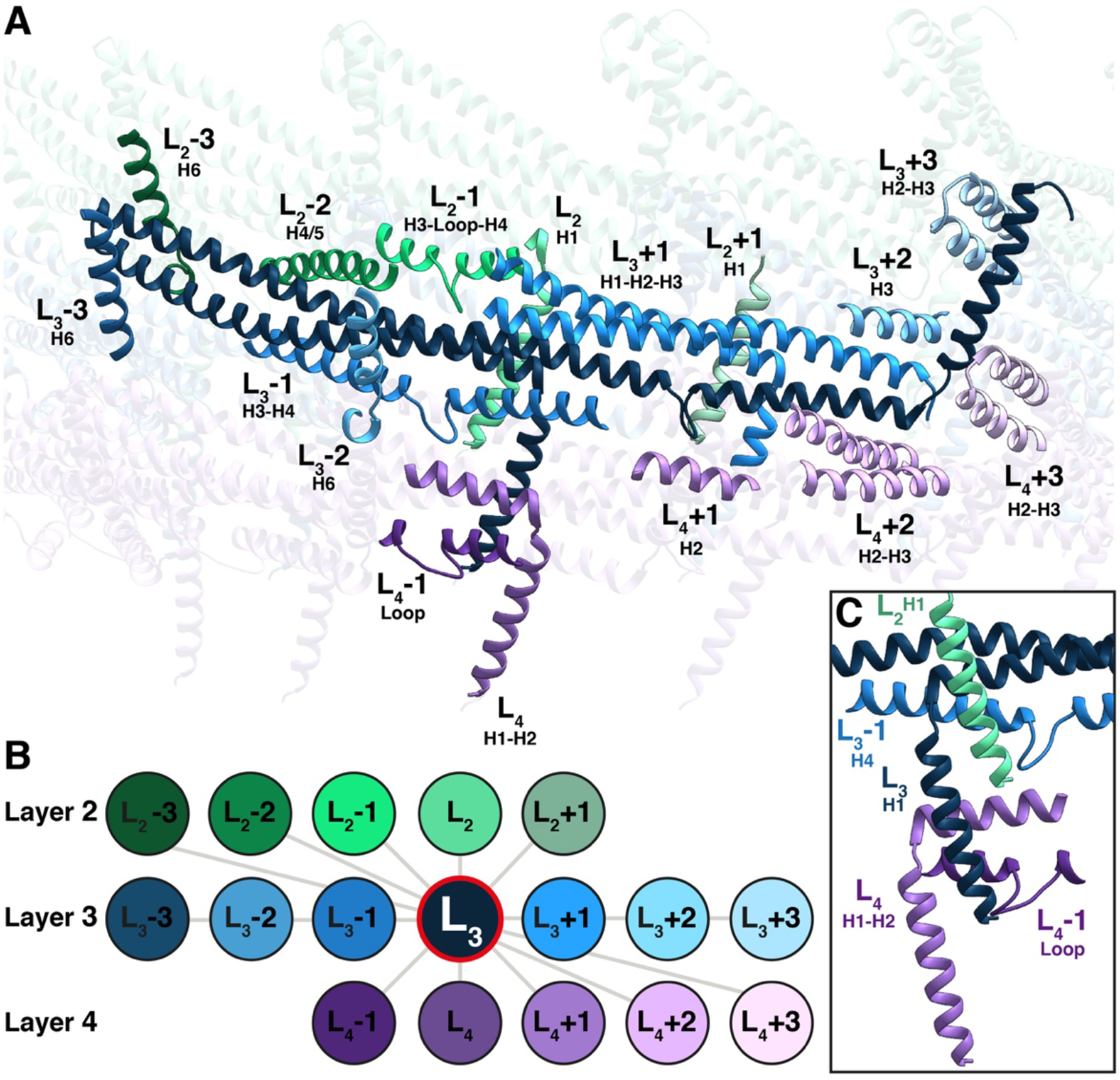
Network of interactions for one VIPP1 monomer within the ring. **(A)** One VIPP1 monomer from layer 3 (L_3_, dark blue) was examined to highlight the intricate network of interactions in the VIPP1 ring. Also displayed are regions of other VIPP1 monomers that are in close proximity and can interact with the L_3_ reference monomer. Numbering is assigned with respect to L_3_: monomers that are positioned to the right of L_3_ are labeled +1, +2 and +3, whereas monomers that are positioned to the left of L_3_ are labeled −1,−2 and −3. Monomers that are directly above and below L_3_ are labeled L_2_ and L_4_, as these monomers come from layer 2 and layer 4, respectively. **(B)** The schematic diagram of this interaction network, showing that one monomer can interact with up to 16 neighbors. **(C)** N-terminal H1 interacts with four other VIPP1 monomers: one monomer from the layer above (layer 2, in this case), one monomer from within the same layer (layer 3, in this case) and two monomers from the layer below (layer 4, in this case).

**Figure S6.**
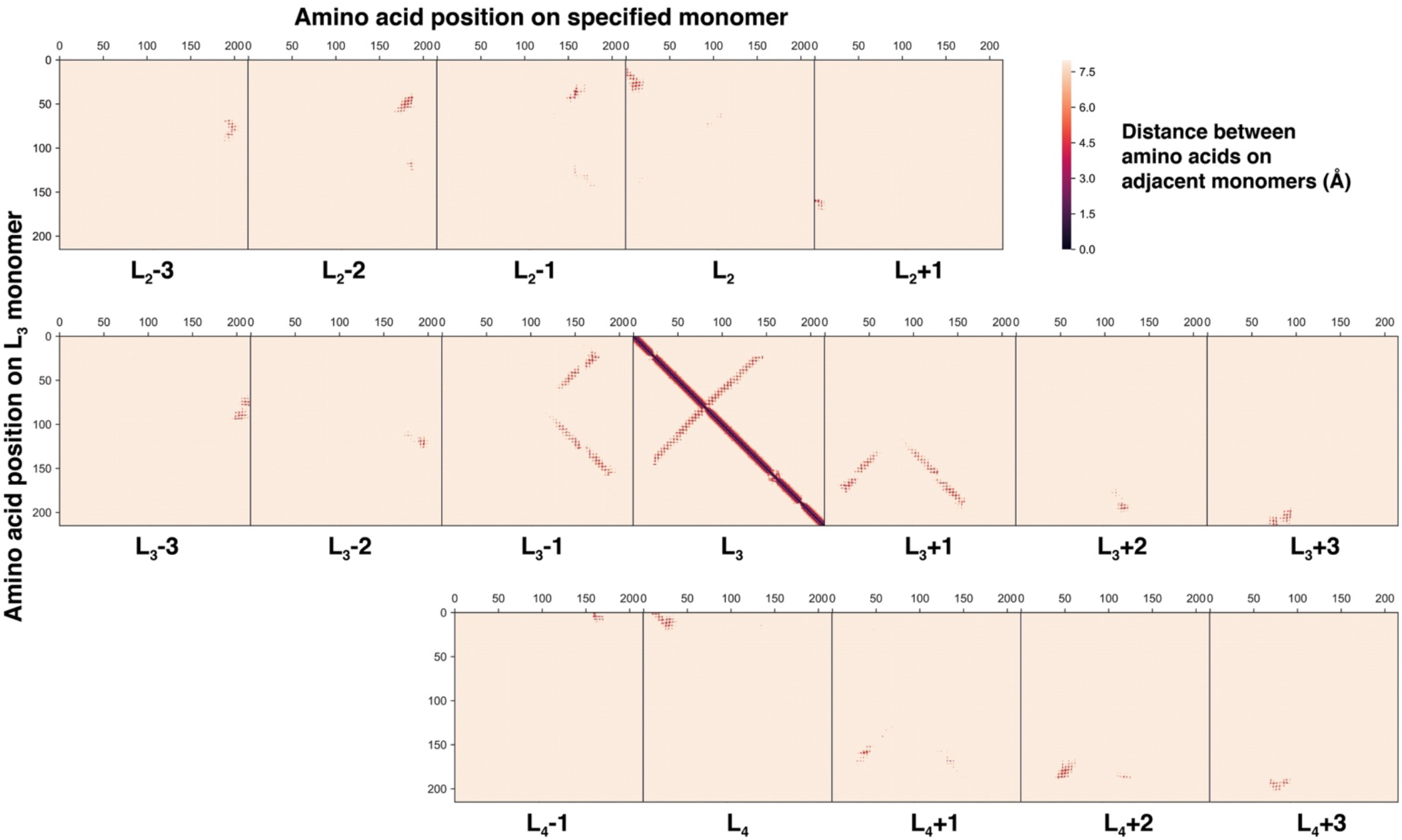
Interaction map for one VIPP1 monomer within the ring. The monomer naming corresponds to Fig. S5. Each square plots the shortest distance between each amino acid residue in the L_3_ monomer (Y-axis) to each amino acid in an adjacent monomer from layers 2, 3, and 4 (X-axis). Distances are color-coded according to the scale bar, with red colors indicating distances <6Å. Patches of red represent interaction interfaces between the monomers. The solid diagonal line from the top left to the bottom right of the central L_3_ x L_3_ square is autocorrelation. The second, lighter diagonal line in this square is the hairpin coiled-coil interaction between VIPP1 monomer’s H2 and H3 domains (see Fig. 1G).

**Figure S7.**
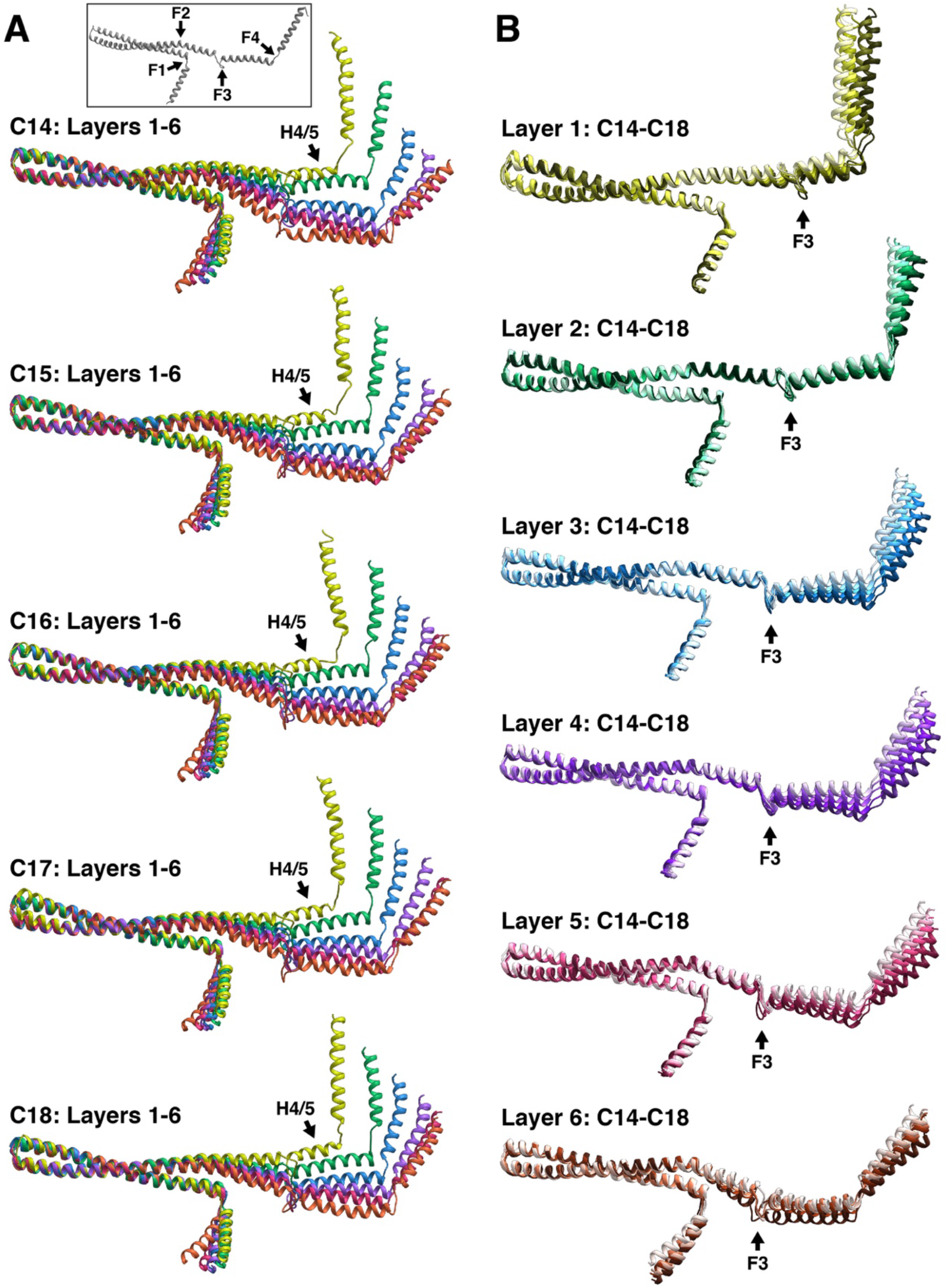
VIPP1 monomer flexibility across ring layers and symmetries. Accompanies Fig. 1I-K. **(A, inset)** VIPP1 monomers have four regions of flexibility (F1-F4). **(A)**All four regions flex to build a ring: VIPP1 monomers from each layer superimposed for rings of each symmetry. In layer 1, part of H4/5 is remodeled into a coil (see Fig. S8). The color scheme matches Figs. 1 and S4. **(B)** Only F3 flexes to accommodate different symmetries: superposition of VIPP1 monomers from the same layers across all five symmetries (five shades of color, with C14 the darkest and C18 the lightest).

**Figure S8.**
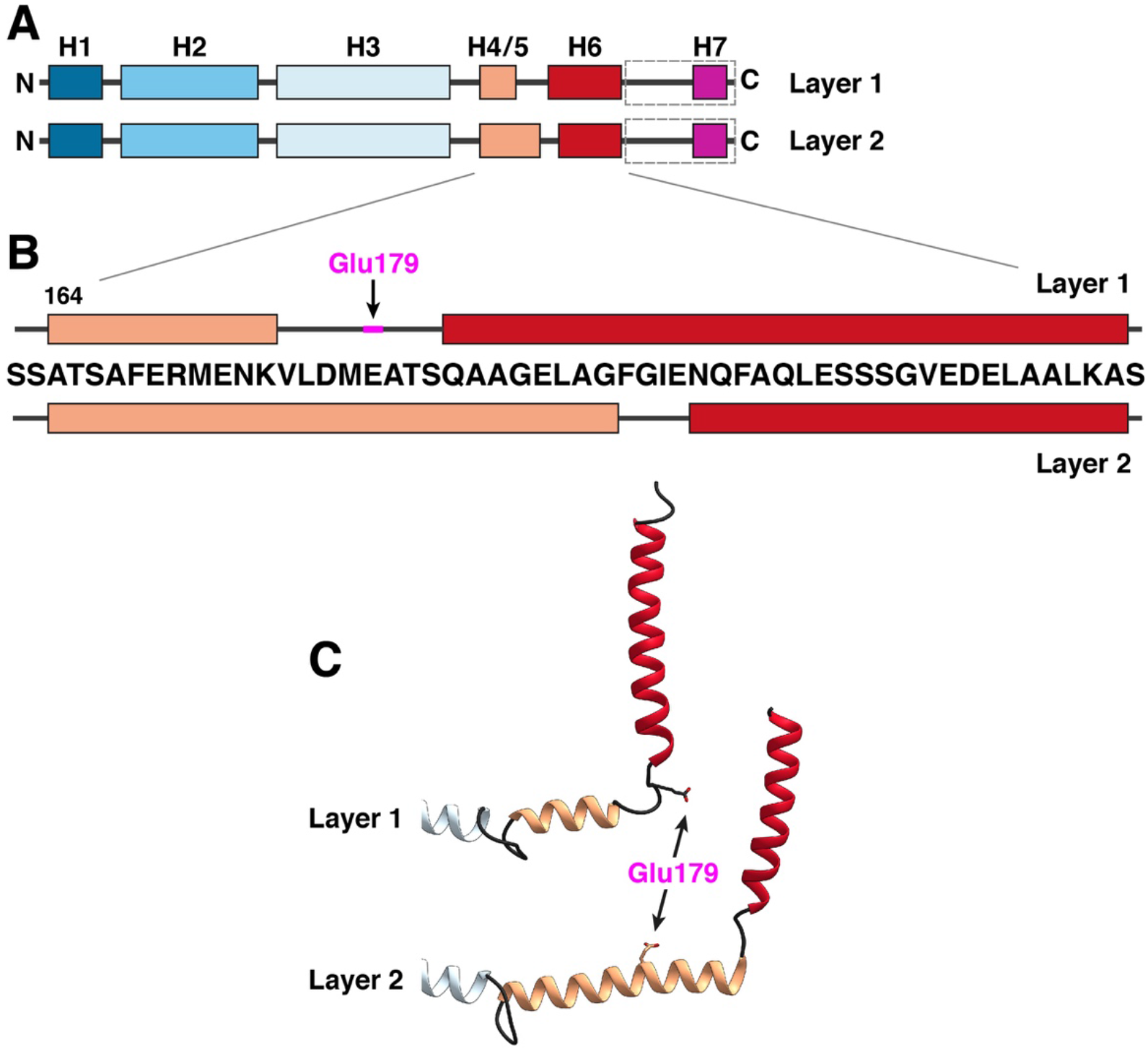
Helix to coil transition in layer 1 helps form the nucleotide binding site. **(A)** Schematic diagram of the VIPP1 monomer’s secondary structure elements (helices H1-H7 in colored rectangles), as shown in Fig. 1F. **(B-C)** Zoom in on the amino acid sequence spanning H4/5 and H6, diagramming the remodeling these two helices between layers 1 and 2. **(C)** Corresponding view of the molecular model, showing the remodeling that occurs between layers 1 and 2. Glu179, which plays a key role in the nucleotide binding pocket, is indicated in magenta. Notice that Glu179 is part of the H4/5 helix in layer 2. However, H4/5 partially opens into a coil in layer 1, placing Glu179 in the loop between the helices as part of the nucleotide binding pocket.

**Figure S9.**
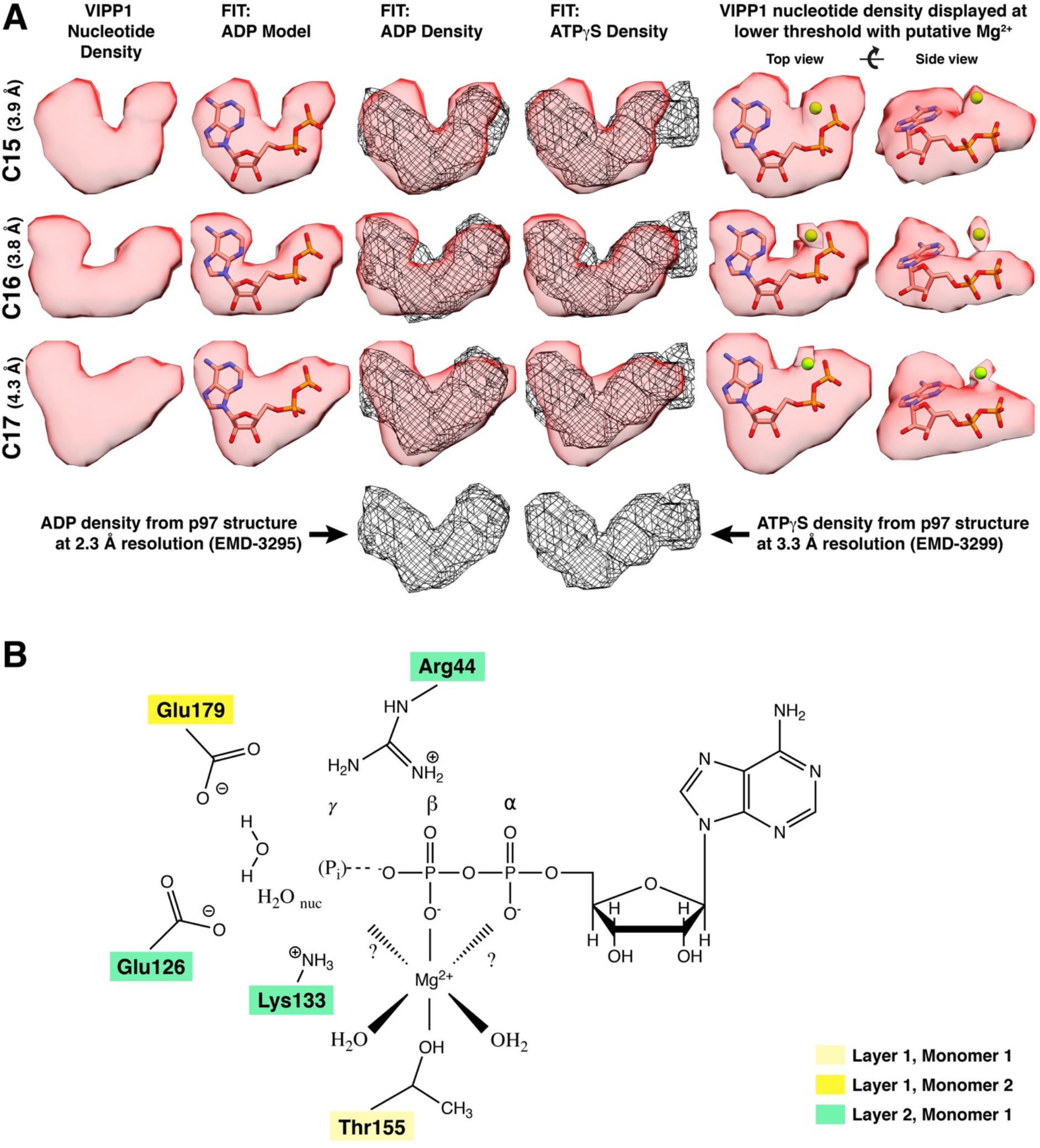
Analysis of the VIPP1 nucleotide density and scheme of the binding pocket. **(A)** Structural comparison of the EM densities corresponding to putative nucleotides observed in VIPP1 maps of three different symmetries (C15, C15, C17; first column from the left) with ADP structural models and cryo-EM densities from known nucleotides. Second column: VIPP1 densities fit with an ADP model. Third and fourth column: VIPP1 densities fit with cryo-EM density from structures of the p97 complex (Banerjee et al., 2016) known to correspond to ADP and ATP*γ*S (black mesh). Notice that the ADP density is a better fit. Fifth and sixth columns: VIPP1 densities displayed at a lower threshold, which grows the occupied volume and reveals a small additional density that may correspond to Mg^2+^. The VIPP1 densities are fit with a structural model of ADP with an Mg^2+^ (green sphere). **(B)** Proposed model for coordination of the nucleotide, Mg^2+^, and a water molecule in the binding pocket. Residues are color coded according to layer and monomer, in accordance with Fig. 2C-E.

**Figure S10.**
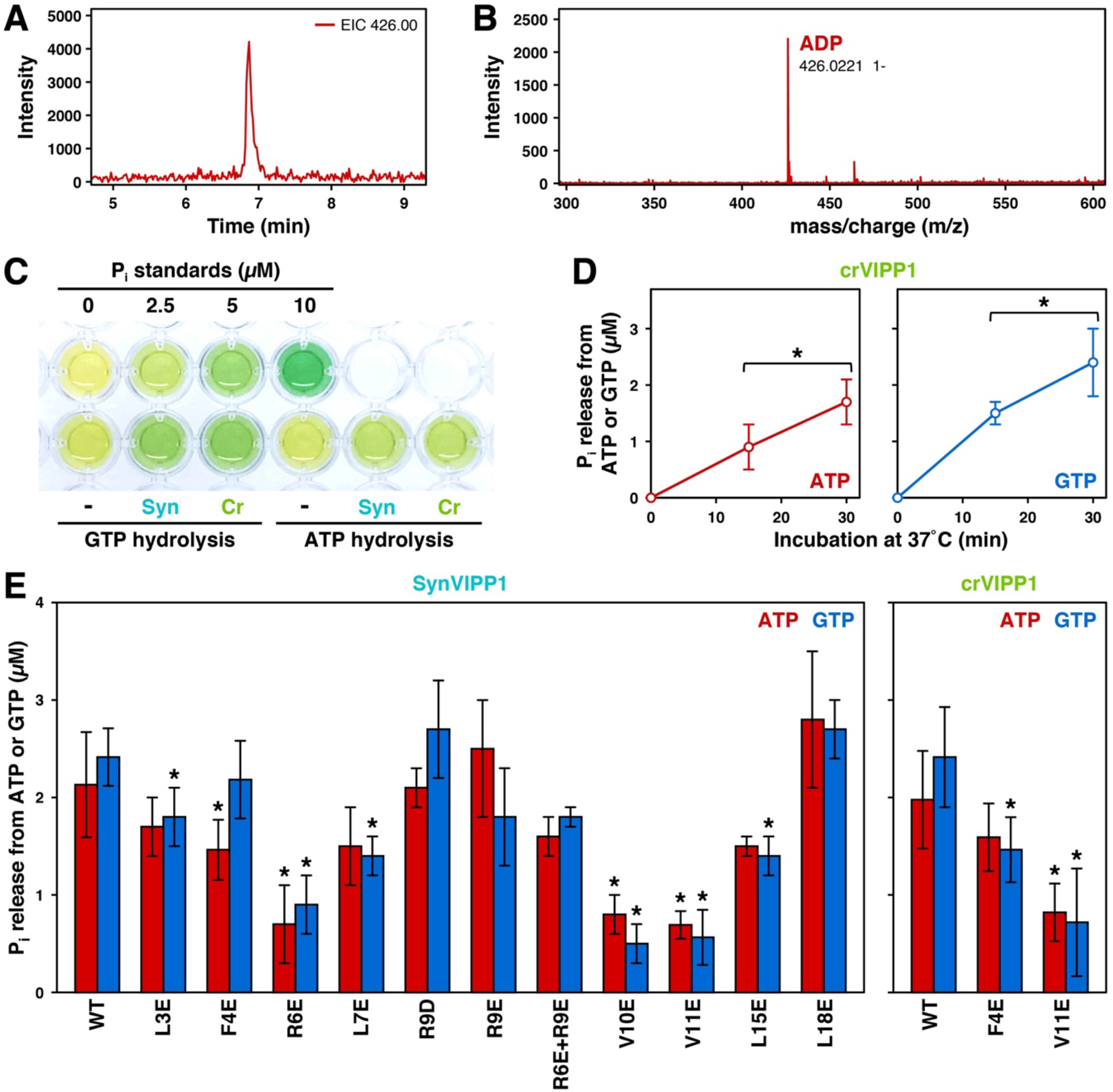
Supplemental mass spectrometry and nucleotide hydrolysis activity data. **(A)** RPIP-HPLC profiles and **(B)**subsequent ESI-MS of isolated wild-type *syn*VIPP1. A distinct mass for ADP was observed. **(C)**Image of a microwell plate containing an *in vitro* nucleotide hydrolysis assay. Inorganic phosphate (P_i_) standards are arrayed in the top row, and the hydrolysis of ATP and GTP by *syn*VIPP1 (Syn) and *cr*VIPP1 (Cr) are assayed on the bottom row (control without protein indicated with “-”). **(D)**Hydrolysis of ATP and GTP by *cr*VIPP1is linear as a function of time. Comparable *syn*VIPP1 measurements are shown in Fig. 2G. **(E)** *In vitro* ATP and GTP hydrolysis by wild-type *syn*VIPP1 (left panel) and *cr*VIPP1 (right panel) compared to point mutations in the N-terminal H1 helix. Error bars are standard deviation from 3-8 replicates, and asterisks indicate a significant change (p<0.05, Welch’s t-test).

**Figure S11.**
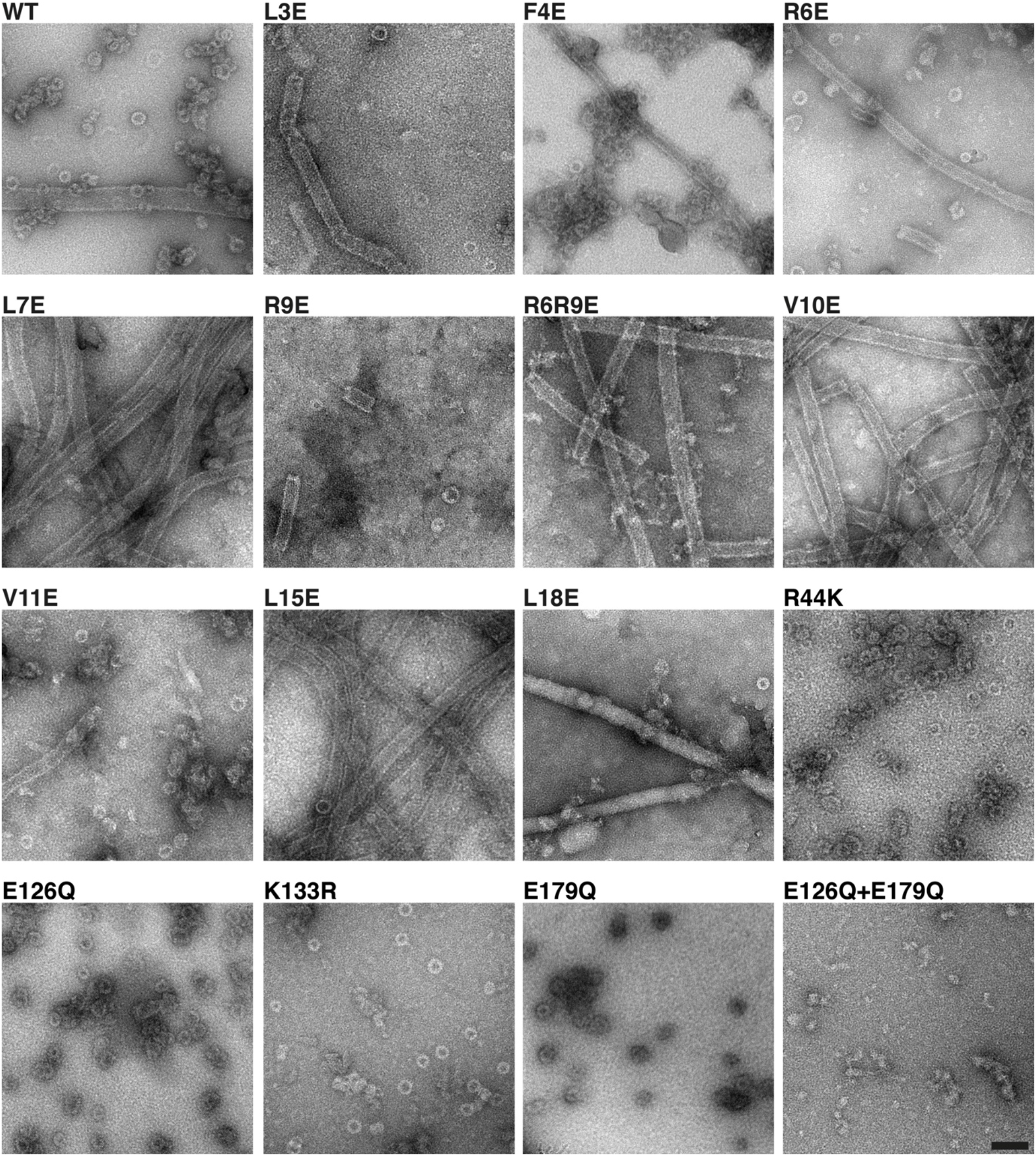
Example negative stain images from each *syn* VIPP1 protein. Mutant names are indicated above each image. While rods were commonly formed by mutations in the H1 helix (L3E through L18E), mutation of nucleotide pocket residues (R44K through E179Q) yielded a WT-like distribution of rings. The E126Q+E179Q double mutant contained abundant small fragments and larger protein aggregates, but no ring or rod structures. Rods were occasionally observed in the WT sample and seemed to be more common at higher protein concentrations. Quantification of the abundance of rings and rods is shown in Fig. 3F. Scale bar: 100 nm.

**Figure S12.**
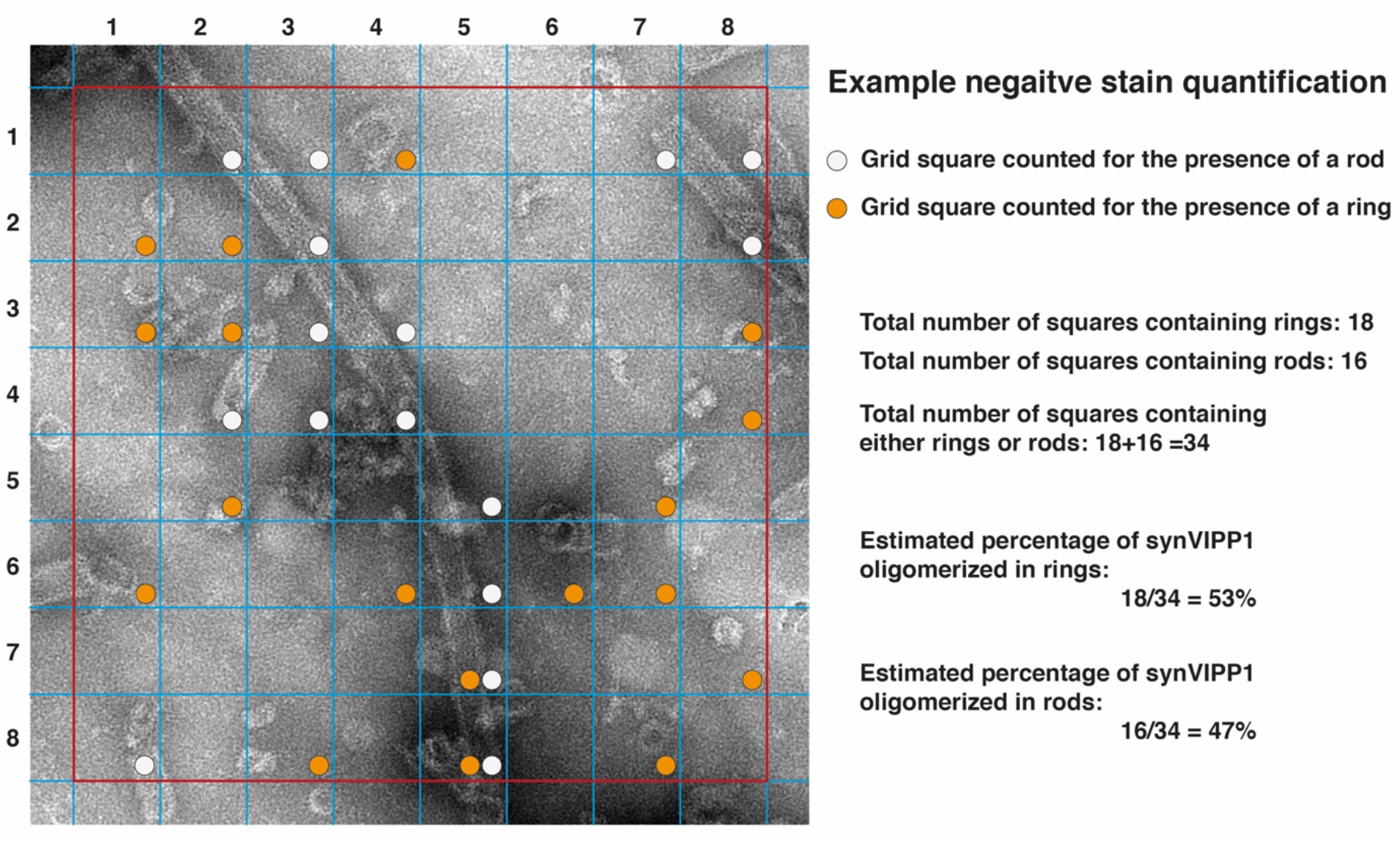
Estimation of the percentage of VIPP1 protein assembled into rings vs. rods. Accompanies the results in Fig. 3F. Negative stain EM images were acquired for wild-type synVIPP1 protein and each of the H1 point mutants (Fig. S11). The images were overlaid with an 8×8 grid map (the grid map border outlined in red; individual 80×80 Å grid squares outlined in blue). Grid squares were manually scored for the presence of rings (orange circles) and/or rods (white circles). If more than 20-30% of a structure was contained within a grid square, that square was counted. The percentage of VIPP1 oligomerized into rings was roughly estimated by dividing the number of grid squares containing rings by the sum of the squares containing rings and squares containing rods (see calculation on the right). For each measurement in Fig. 3F, we analyzed 20-50 images from 2-3 grids, produced from 2-4 independent protein preps.

**Figure S13.**
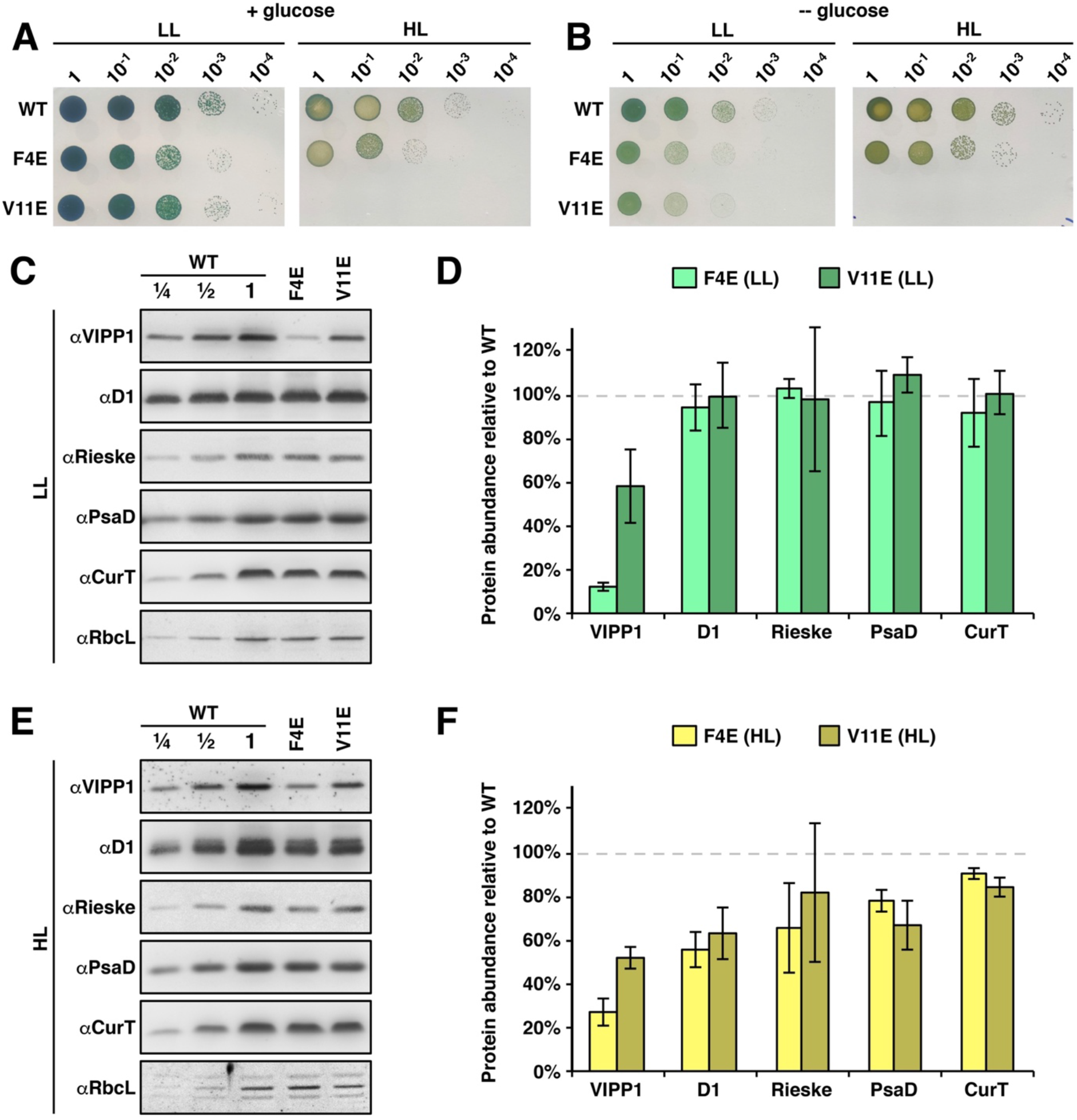
Supplementary data for the *in vivo* H1 point mutants. **(A-B)** Spot growth tests of WT (control where endogenous VIPP1 was replaced by the wild-type gene), F4E, and V11E stains after five days on agar plates in low light (LL, 30 μmol photons m^−2^s^−1^) and high light (HL, 200 μmol photons m^−2^s^−1^). **(A)**Cells grown on plates containing glucose. These images are reproduced from Fig. 4A but displayed with more natural contrast. **(B)** Cells grown on plates without glucose. **(C, E)** Western blot analysis of various thylakoid proteins in WT, F4E, and V11E in LL (panel C) and after switch to HL for 24 hours (panel E). The Rubisco large subunit RbcL was used as a loading control. **(D, F)** Relative changes of protein levels in the F4E an V11E mutants compared to WT, quantified from the Western blots in panels C and E. Plots show mean and standard deviation (error bars) from three independent biological replicates.

**Figure S14.**
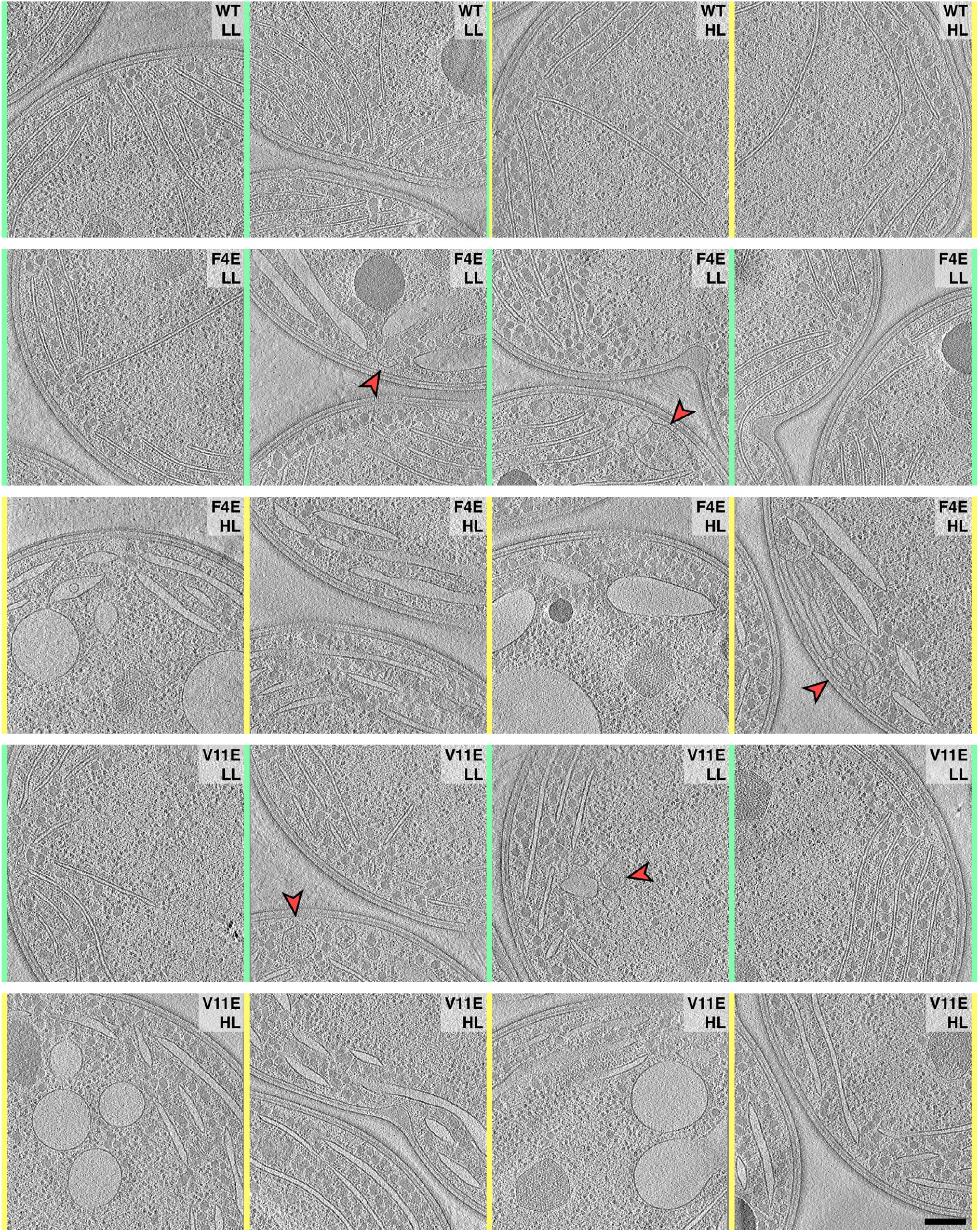
Tomogram gallery. Overviews of additional tomograms from the WT, F4E, and V11E strains grown in low light (LL, green) and after 24 hours in high light (HL, yellow). Quantification of luminal widths is shown in Fig. 4L. Red arrowhead indicates abnormal convergence zone architecture (more examples in Fig. 5). Scale bar: 200 nm.

## Supplementary Movies

**Movie 1.** Tour of the C15 *syn*VIPP1 cryo-EM density map. The movie tilts from top view to side view, then shows how monomers assemble into layers and layers assemble into the full ring structure. The ring is then sliced open to reveal the luminal columns of H1 helices. The extra densities corresponding to ADP nucleotide (red) and H1-bound lipids (orange) are then revealed. Accompanies Figs. 1–3.

**Movie 2.** Morph map of the VIPP1 model showing monomer flexibility within the C16 ring. Accompanies Fig. 1J.

**Movie 3.** Morph map of the VIPP1 model showing the flexibility of a layer 3 monomer from C14 to C18. Accompanies Fig. 1K.

**Movie 4.** *In situ* cryo-ET of a WT *Synechocystis* cell grown for 24 hours in high light. Sequential sections back and forth through the tomographic volume shown in Fig. 4D, followed by reveal of the 3D segmentation from Fig. 4E (orthographic view).

**Movie 5.** *In situ* cryo-ET of an F4E *Synechocystis* cell grown for 24 hours in high light. Sequential sections back and forth through the tomographic volume shown in Figs. 4G and 5A, followed by reveal of the 3D segmentation from Figs. 4H and 5B-C (orthographic view). Notice the aberrant membrane structures at the thylakoid convergence zone.

**Movie 6.** *In situ* cryo-ET of a V11E *Synechocystis* cell grown for 24 hours in high light. Sequential sections back and forth through the tomographic volume shown in Fig. 4J, followed by reveal of the 3D segmentation from Fig. 4K (orthographic view).

## Methods

### *syn* VIPP1 vector design, protein expression and purification

The coding region of wild-type *syn*VIPP1 was amplified from *Synechocystis* genomic DNA using primers VIPPSyn-Spe-Sap and VIPPSyn-Xho. The 839-bp PCR product was digested with SapI and XhoI and ligated into SapI–XhoI-digested pTYB11 (NEB), generating pMS451. Point mutants were introduced by PCR (30 sec at 98°C, 20 sec at 63-70°C, 7.5 min at 72°C) using the primer pairs given in Supplemental Table 1 and pMS451 as template. For the E126Q+E179Q double mutant, the plasmid harboring the E126Q mutant was used as template. PCR products were phosphorylated with polynucleotide kinase (NEB) and circularized with T4 DNA ligase (NEB). Correct cloning was verified by Sanger sequencing. All *syn*VIPP1 variants and the CGE1a protein (pMS300) (Willmund et al., 2007) were produced in *E. coli* ER2566 as C-terminal fusions to a chitin-binding domain/intein and purified by chitin-affinity chromatography. For this, a 10-mL overnight culture was diluted into 1 L TB containing 100 μg/mL Ampicillin and grown at 37°C for 7-8 h. IPTG was then added to a final concentration of 0.5 mM and growth continued at 18°C overnight. Cells were harvested by centrifugation at 5,000 g and 4°C for 10 min, resuspended in 25 ml ice-cold lysis buffer (20 mM Hepes-KOH pH 8.0, 0.5 M NaCl, 1 mM EDTA, 0.1% Triton X-100, Roche cOmplete^™^ EDTA-free protease inhibitor cocktail) and sonicated. After centrifugation at 20,000 g and 4°C for 30 min, the supernatant was passed twice at a flow rate of 0.5 mL/min through a column with 6 mL chitin beads (NEB) equilibrated with lysis buffer. The column was then washed with 100 mL lysis buffer at 2 mL/min, followed by a wash with 10 mL KMH buffer (20 mM Hepes-KOH pH 7.6, 80 mM KCl, 2.5 mM MgCl_2_) containing 5 mM ATP to remove DnaK binding to VIPP1. For the GTP-exchange mass spectrometry experiment (Fig. 2F), an additional wash was performed with 5 mM GTP in 10 mL KMH buffer. After a final wash with 20 mL lysis buffer lacking Triton at 2 mL/min, the column was flushed with 10 mL cleavage buffer (20 mM Tris-HCl pH 9.0, 0.5 M NaCl, 1 mM EDTA, 50 mM DTT), gently agitated overnight at room temperature, and *syn*VIPP1 was slowly eluted. Elution was finalized with 5 mL KMH buffer to yield ~10 mL of eluate, which was then concentrated to ~5 mL by centrifugation at 4500 g using a Millipore concentrator (AMICON MWCO 3,000). The concentrate was diluted twice with 20 mL dialysis buffer 1 (20 mM Tris-HCl, pH 7.5, 200 mM NaCl, 75 mM NaSCN) after concentration to 1-2 mL, and diluted twice more with 20 mL dialysis buffer 2 (20 mM Tris-HCl, pH 7.5, 50 mM NaCl, 75 mM NaSCN) after concentration to 1-2 mL. After the final run, the protein concentration was determined by Bradford assay. The purity of the protein preps was confirmed by SDS PAGE followed by Coomassie stain (Fig. S2A). Proteins were either kept on ice for analysis by cryo-EM, or quick-frozen in liquid nitrogen and stored at –80°C.

**Supplemental Table 1.**
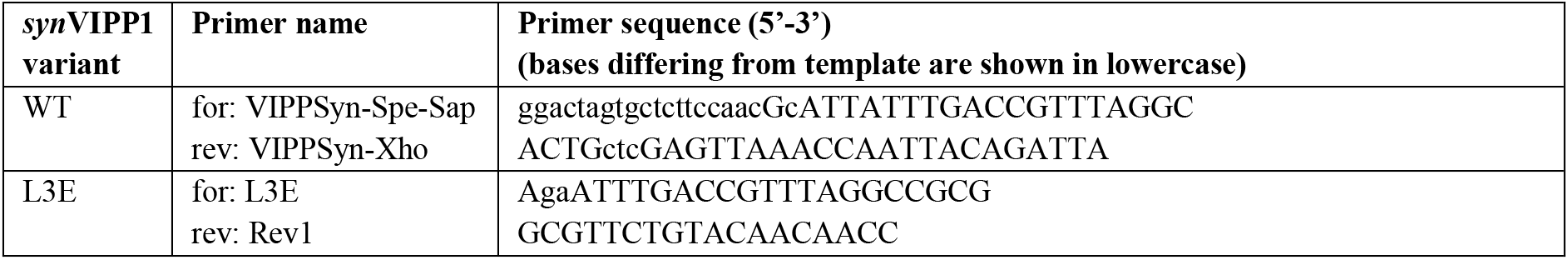
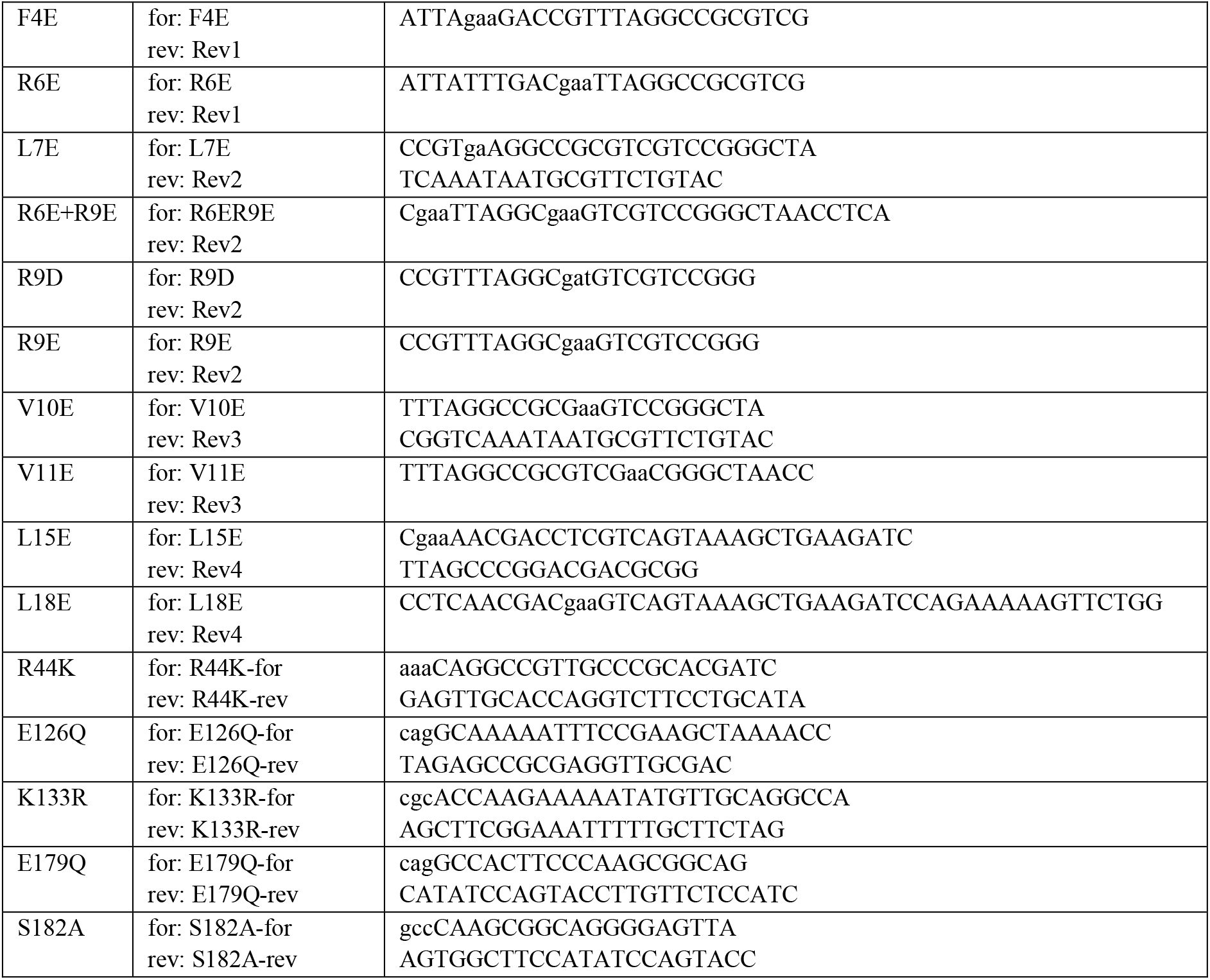
Primers used for amplifying the coding sequence of wild-type *syn*VIPP1 and for site-directed mutagenesis.

### Negative stain EM data acquisition

5 μL of purified synVIPP1 protein was applied to glow discharged, 200 mesh copper grids (G2200C, Plano GmbH) that had been coated with homemade carbon film. The sample was incubated for 2 min, blotted, washed three times with water, and then stained with 2% uranyl acetate for 30 sec. Images were recorded using a Tecnai F20 FEG microscope (FEI, Thermo Fisher Scientific) operated at 200 kV, with an FEI Eagle CCD camera, a calibrated pixel size of 2.21 Å per pixel, and a defocus range of −2 to −5 μm.

### Cryo-EM data acquisition

Cryo-EM grids were prepared with freshly purified synVIPP1 protein. 4 μL of the protein was applied to glow discharged, holey carbon-coated copper grids (R 2/1, 200 mesh, Quantifoil Micro Tools). Grids were plunge-frozen in a liquid ethane/propane mixture using a Vitrobot Mark 4 (FEI, Thermo Fisher Scientific). Blotting chamber conditions were set to 4 °C and 95% humidity, and grids were blotted using blot force 10 and a blot time of 10 sec. Grids were stored in liquid nitrogen until usage. Images were acquired on a Titan Krios microscope (FEI, Thermo Fisher Scientific) at 300 kV, using a post-column energy filter (Quantum, Gatan) and a K2 Summit direct electron detector (Gatan) operated in counting mode. For each image, 80 frames were acquired over 8 sec, with a total dose of ~45 e^−^/Å^2^ and a calibrated pixel size of 1.35 Å. A total of 8120 images were recorded, using a defocus range of −0.5 to −3.5 μm.

### Cryo-EM image processing and analysis

See the workflow in Fig. S1. The frame stacks were subjected to alignment with 5×5 patches and dose filtering using MotionCor2 (Zheng et al., 2017), yielding 8120 aligned and dose-weighted micrograph images (Figs. S1A, S2B). Automated particle picking was performed with crYOLO (Wagner et al., 2019) by training on a small dataset of 80 manually-picked images and then using this trained network to pick particles from the full dataset. All subsequent image processing was carried out using the RELION 3.0 software suite (Zivanov et al., 2018). CTF parameters were determined using Gctf (Zhang, 2016). Based on the crYOLO picking, ~337,000 particles were extracted (defined as Dataset-A). After particle extraction, the dataset was cleaned using reference-free 2D classification (Fig. S2C), yielding ~263,000 particles (defined as Dataset-B).

To generate a reference-free initial 3D model, we used a RELION implementation of stochastic gradient descent (SGD)(Punjani et al., 2017) without applying symmetry (Fig. S1B). The SGD algorithm allows for *ab initio* heterogeneous structure determination without a prior reference map. Starting with Dataset-B, a very rough class was obtained from ~33,000 particles. Another round of SGD was performed using these particles, and ~20,000 particles were selected to produce an improved 3D map exhibiting C17 symmetry. Using this volume as a reference, 3D classification of the ~20,000 particles was performed without applying symmetry constraints. The 3D classification split the particles into several classes, yielding 3D maps for both C15 and C17 symmetries. Using these two volumes as references, a new 3D classification of the full Dataset-B was performed, while enforcing symmetry constraints (Fig. S1C). This split the particles into two large classes (C15 and C17) and generated improved 3D maps to be used as references for the next round of 3D classification. To obtain homogeneous classes of particles, ~120,000 particles from the C15 class and ~62,000 particles from the C17 class were subjected to several rounds of 3D classification (Fig. S1D). These classifications split the particles into new classes with C14, C15, C16, C17 and C18 symmetries. Several rounds of 3D refinement were performed for these symmetry classes, yielding 3D structures with resolutions ranging from 4.5 Å to 6.5 Å (Fig. S1E).

When further rounds of 3D classification and refinement did not improve the resolution of Dataset-B, the final density maps of each symmetry were used as references for 3D classification of the full Dataset-A (~337,000 particles) (Fig. S1F-G). The reason behind this was to extract the particles directly from the initial dataset based on the specific 3D references corresponding to each symmetry class. Coarse angular sampling (7.5°) was applied to speed up the classification process, and the dataset was divided into five classes corresponding to specific symmetries (Fig. S1F). Each symmetry class was then subjected to several rounds of 3D classification with fine angular sampling (3.5°) (Fig. S1G). The goal of this step was to clean the classes further, increasing the structural homogeneity of the particles.

The final classes were next processed through iterative rounds of RELION 3D auto-refinement to improve the alignment, per-particle CTF refinement to estimate the defocus and astigmatism of each particle, and Bayesian polishing to correct for the motion of each particle (using frames 3-50) and filter for the radiation damage. Alignment was carried out following the “gold-standard” procedure, which randomly splits the dataset into two half sets and aligns them independently. This allowed calculation of a Fourier shell correlation (FSC) curve for resolution estimation (Fig. S3A) using the FSC=0.143 threshold criterion (Rosenthal and Henderson, 2003). Local resolution maps (Fig. S3B) were generated with the RELION local resolution function (Cardone et al., 2013). Images of the cryo-EM density maps were generated with UCSF Chimera (Pettersen et al., 2004) and ChimeraX (Goddard et al., 2018).

### Modeling of VIPP1 into the cryo-EM density

The initial PDB structure of *syn*VIPP1 was obtained by homology modeling with Modeler software (Sali and Blundell, 1993) starting with the available crystal structure of *E. coli* PspA (PDB: 4WHE, corresponding to helices H2 and H3 of VIPP1) (Osadnik et al., 2015) and the *syn*VIPP1 sequence (UniProt: A0A068MW27). Domains not resolved in the crystal structure (residues 2-25 and 142-216, corresponding to helices H1, H4/5, and H6) were modeled using the Rosetta (Leaver-Fay et al., 2011) *ab initio* structure prediction framework implemented as plugin in VMD software (Humphrey et al., 1996). This framework was initially developed to furnish the structurally unresolved regions of the 26S proteasome (Schweitzer et al., 2016). We did not model the VIPP1 C-terminus (residues 217-267, corresponding to H7 and a disordered linker), as it was not resolved within the EM density map, presumably due to its highly dynamic nature (Fig. 1F-G). The models were clustered using the k-means algorithm. For each symmetry, single subunits for each layer of the VIPP1 ring were segmented.

Each of the clusters obtained was fitted into the segmented maps using molecular dynamics flexible fitting (MDFF) (Trabuco et al., 2008). MDFF employs molecular dynamics to fit initial models into a density in real space, and thus, permits protein flexibility while maintaining realistic protein conformations (Goh et al., 2016). We used NAMD (Phillips et al., 2005) with the CHARMM36 force field for MDFF calculations. During MDFF runs, restraints to preserve the secondary structure, chirality, and cis-peptide bonds were applied to avoid overfitting. As further step to reduce artifacts due to overfitting all MDFF runs were performed at a modest gscale of 0.3.

After MDFF convergence, the cross-correlation coefficient for the clusters was calculated, and the structure showing the highest correlation was used for subsequent steps. Once subunits for each segment were fitted by interactive MDFF, the entire structure was created by applying the respective symmetry operation (C14, C15, C16, C17, and C18) on the model. The whole structure was then refined using MDFF and symmetry-adapted constraints to apply a harmonic potential to the symmetry-averaged positions of each individual VIPP1 subunit within the complete ring. After this refinement, the structures were re-symmetrized with Biopython (Hamelryck and Manderick, 2003) using six selected monomers, one from each layer of the VIPP1 ring. Steric clashes between the symmetry subunits were removed by MDFF energy minimization of the whole structure into the cryo-EM density map. The interaction network between VIPP1 monomers in the ring (Fig. S6) was generated using MDAnalysis (Michaud-Agrawal et al., 2011) based on the distances between residues within the VIPP1 structure.

The initial prediction of a nucleotide binding site in *syn*VIPP1 was performed using the GTPbinder (Chauhan et al., 2010) and ATPint (Chauhan et al., 2009) web servers. AutoDock4 (Morris et al., 2009) was used to predict possible nucleotide binding modes. We aligned the ADP from the human 26S proteasome structure (PDB: 5L4G) (Schweitzer et al., 2016) with the best scoring result. The model of the nucleotide binding region was further refined to the density in real space by an iterative combination of MDFF with *ab initio* structure prediction as well as Monte Carlo-based backbone and side-chain rotamer search algorithms, following the strategy detailed in (Wehmer et al., 2017). Subsequently, we further refined the nucleotide binding pocket using interactive MDFF to manually pull side chains into EM density, while interactively checking the cross-correlation values in VMD.

Finally, the complete structure was refined with Phenix software (Liebschner et al., 2019) real space refinement with reference coordinate restraints (σ=0.05) on the nucleotide binding site. The resulting structure was re-symmetrized.

**Supplemental Table 2.** Cryo-EM map and molecular model statistics.

| Structure | C14 | C15 | C16 | C17 | C18 |
| --- | --- | --- | --- | --- | --- |
| Model resolution (Å) | 4.9 | 3.9 | 3.8 | 4.3 | 5.0 |
| FSC threshold | 0.143 | 0.143 | 0.143 | 0.143 | 0.143 |
| Map sharpening B factor (Å<sup>2</sup>) | -164.37 | -136.05 | -154.73 | -162.25 | -214.50 |
| **Model composition** |  |  |  |  |  |
| Non-hydrogen atoms | 9879 | 9887 | 9887 | 9887 | 9887 |
| Protein atoms | 9852 | 9860 | 9860 | 9860 | 9860 |
| Ligand atoms | 27 | 27 | 27 | 27 | 27 |
| **Rms deviation** |  |  |  |  |  |
| Bond length (Å) | 0.017 | 0.018 | 0.018 | 0.018 | 0.018 |
| Bond angles (°) | 1.656 | 1.783 | 1.274 | 1.738 | 1.708 |
| **Validation** |  |  |  |  |  |
| Molprobity Score | 1.59 | 1.23 | 1.29 | 1.33 | 1.80 |
| Clash score | 7.03 | 4.45 | 4.05 | 3.95 | 6.27 |
| Poor rotamers (%) | 0.4 | 0.51 | 1.11 | 0.91 | 1.41 |
| **Ramachandran plot** |  |  |  |  |  |
| Favored (%) | 96.78 | 97.96 | 97.72 | 97.71 | 95.05 |
| Allowed (%) | 3.07 | 1.49 | 2.28 | 2.20 | 4.08 |
| Disallowed (%) | 0.16 | 0.55 | 0 | 0.63 | 0.86 |

### Detection by VIPP1-bound nucleotide by mass spectrometry

The ATP and GTP wash steps were performed during purification while the VIPP1 protein was bound to the chitin column, as described above. Purified VIPP1 at a concentration of 1 μM was injected onto a C18 column (Agilent Zorbax Eclipse Plus C18, 2.1 × 100 mm2, pore size 3.5 μm), mounted on an Agilent 1290 HPLC apparatus. Bound ligands were eluted at 250 μL/min using a gradient of buffer A (20 mM triethylammonium acetate (pH 9.0)) and buffer B (20 mM triethylammonium acetate (pH 9.0), 20% acetonitrile) as previously described (Gat et al., 2019). The experiment was conducted at pH 9.0, with the mass spectrometer (maXis II ETD, Bruker) operating in negative ion mode, with a mass range of 200–1,200 m/z and collision cell energy of −10.0 eV to accommodate detection of the negatively charged compounds. Therefore, the masses of both ADP and GDP are off by −1 Da (Figs. 2F, S10A-B).

### VIPP1 nucleotide hydrolysis assays

ATP and GTP hydrolysis assays were carried out as previously described (Ohnishi et al., 2018) with slight modifications, using ATPase and GTPase assay kits (Expedeon Ltd.) that employ a malachite green-based detection system with PiColorLock^™^. All reactions were performed in a 200 μL volume according to the manufacturer instructions. The reaction mixture contained 50 mM Tris-HCl (pH 7.5), 2.5 mM MgCl_2_, 0.5 mM ATP or GTP, and purified *syn*VIPP1 or *cr*VIPP1 recombinant protein. The ATP hydrolysis assay was performed with 12 μg of recombinant protein, whereas the GTP hydrolysis assay was performed with either 6 μg of *syn*VIPP1 or 4 μg of *cr*VIPP1 protein. Reaction mixtures were incubated at 37 °C for 30 min and were then immediately cooled in ice water for 1 min. Subsequently, each reaction was terminated by adding 50 μL of PiColorLock^™^ at room temperature. Unless specifically described, the 30 min incubation time was used as the standard condition. The released Pi in each reaction mixture was quantified by measuring absorption at 635 nm according to the calibration curve obtained from the absorption of Pi standard solutions (see Fig. S10C). Because stock solutions of ATP and GTP always contain a trace amount of free Pi, measurements were normalized by the background absorption value of reaction mixture that had not been supplemented with recombinant protein to obtain VIPP1-dependent Pi release.

### Estimation of the distribution of rings and rods in *syn*VIPP1 mutant proteins

The point mutations we inserted into N-terminal H1 caused some of *syn*VIPP1 to assemble into rods instead of rings. We performed a grid-based measurement of negative stain EM images to roughly estimate the percentage of VIPP1 protein incorporated within rings and rods for each of the mutants (Fig. S12).

The dimensions of the negative stain micrographs were 4096×4096 pixels, with a pixel size of 0.184 Å. Therefore, each micrograph was a square with a side length of 754 Å (4096 pixels multiplied by 0.184 Å). The diameters of Vipp1 rings and rods were roughly 30-40 nm. Based on the Vipp1 dimensions, a grid spacing of 80 Å was chosen, yielding a grid map of 8×8 squares. The 8×8 grid map was overlaid on each of the images, and the presence of rings and/or rods in each square was manually counted. For both rings and rods, if more than 20-30% of a structure was contained within a grid square, that square was counted. Rods frequently extended through several squares, resulting in multiple counts for one structure. Thus, the goal of this analysis was not to measure the relative abundance of ring and rod structures, but rather to estimate the percentage of Vipp1 protein incorporated into the two types of structures. The cumulative results are shown in Fig. 3F.

### Generation of *Synechocystis* mutant strains

To create the *Synechocystis* mutants as well as a control strain, a construct was designed containing the vipp1 gene with its flanking non-coding regions up to 549 bp upstream and 47 bp downstream of the vipp1 gene. A kanamycin resistance cassette (sequence taken from pBSL14 (ATCC 87127)) was inserted 149 bp upstream of vipp1. The construct was ordered as a gBlock DNA fragment (Integrated DNA Technologies), blunt-cloned in the pJET1.2 vector (Thermo Scientific) and subsequently subjected to site-directed mutagenesis (QuikChange XL Site-Directed Mutagenesis Kit, Agilent Technologies) according to manufacturer’s instructions to introduce the mutations F4E or V11E (primer pairs vipp1-F4E-1 + vipp1-F4E-2 and vipp1-V11E-1 + vipp1-V11E-2, Supplemental Table 3). The sequences of both mutant constructs as well as the control construct were verified by sequencing and transformed into *Synechocystis* wild-type cells as previously described (Eaton-Rye, 2004). Complete segregation of the resulting control and mutant strains was verified by PCR (primers vipp1-seg-fw and vipp1-seg-rev, Supplemental Table 3), and the insertion of the point mutations during homologous recombination was confirmed again by sequencing (primer vipp1-seg-rev, Supplemental Table 3).

**Supplemental Table 3.**
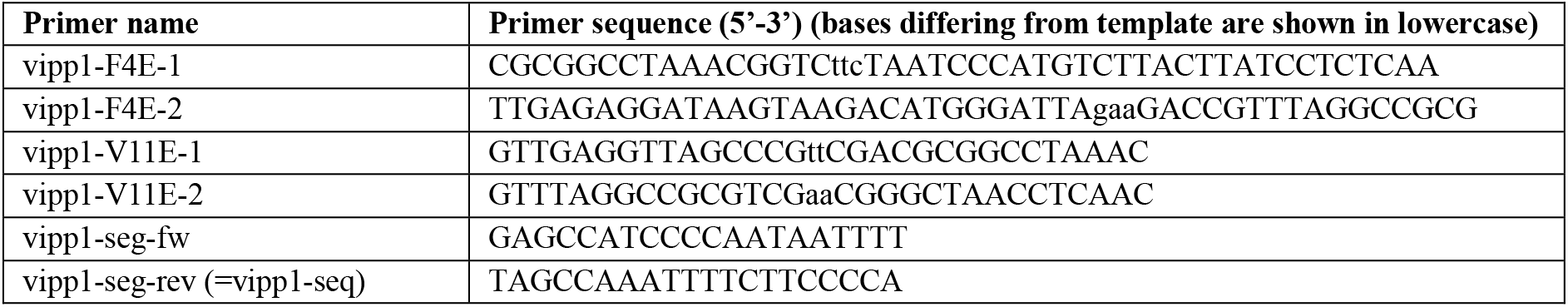
Primers used for generation of *Synechocystis* mutant strains via site-directed mutagenesis

### Physiological and biochemical analysis of *Synechocystis* mutants

*Synechocystis* strains were grown on solid or in liquid BG11 medium (Rippka et al., 1979) at 30 °C and 30 μmol photons m^−2^ s^−1^ in presence of 5 mM glucose if not stated otherwise. For selection of mutant strains, up to 200 μg/ml kanamycin was added to solid BG11 medium. For growth analysis, log-phase *Synechocystis* cultures grown in low light (30 μmol photons m^−2^ s^−1^) were adjusted to OD_750_=1 and dilution series were prepared up to OD_750_=10^−4^. Next, 5 μl of each dilution and strain were spotted on solid BG11 with and without glucose. Plates were incubated in low light and high light (200 μmol photons m^−2^ s^−1^) and the results were documented after 5 days of growth.

For analysis of whole cell protein accumulation, *Synechocystis* pre-cultures were grown in low light for 2 days until they reached log phase. Main cultures were inoculated to OD_750_=0.1 and grown in low light or high light for 24 h. Cells were harvested by centrifugation, resuspended (50 mM Tris-HCl pH 7, 20 mM MgCl_2_, 20 mM KCl, 0.5% Triton X-100) and broken using a BeadBug Homogenizer (Biozym) and glass beads in three cycles of alternating 20 s shaking and 1 min incubation on ice. Lysates were incubated on ice for 15 min, centrifuged (20,000 g, 1 min 4 °C) and the supernatant protein concentrations were determined by Bradford assay using RotiQuant (Carl Roth GmbH + Co. KG). Samples were subjected to Western blot analysis, and quantification of the results from three independent biological replicates was performed using the ImageQuant TL software (GE Healthcare). Generation of the antibodies used in this study has been described previously: αCurT (Heinz et al., 2016), αD1 (Schottkowski et al., 2009) and αVIPP1 (Aseeva et al., 2007). Additional antibodies were purchased from Agrisera: αPsaD, αRbcL, αRieske and αS1.

### *In situ* cryo-ET data acquisition

Growth of *Synechocystis* strains for *in situ* cryo-ET was performed identically as described for Western analysis. Using a Vitrobot Mark 4 (FEI, Thermo Fisher Scientific), 4 μL of cell culture was blotted onto R 1/4 SiO_2_-coated 200-mesh gold EM grids (Quantifoil Micro Tools) and plunge frozen in a liquid ethane/propane mixture. Grids were stored in liquid nitrogen until used for FIB milling. Cryo-FIB milling was performed as previously described (Schaffer et al., 2015; Schaffer et al., 2017), using an Aquilos dual-beam FIB/SEM instrument (FEI, Thermo Fisher Scientific). Grids were clipped into Autogrid support rings modified with a cut-out that allows access to the ion beam at low angle (FEI, Thermo Fisher Scientific). In the FIB/SEM chamber, the grids were sputter-coated with metallic platinum to reduce charging, then coated with a thicker layer of organometallic platinum using a gas injection system to protect the sample surface. The cells were then milled with a gallium ion beam to produce lamellas of 100-200 nm thickness, and the grids were transferred out of the FIB/SEM and stored in liquid nitrogen until imaged by cryo-ET.

Grids containing milled samples were transferred into a 300 kV Titan Krios microscope (FEI Thermo Fisher), equipped with a post-column energy filter (Quantum, Gatan), and a direct detector camera (K2 summit, Gatan). Using SerialEM software (Mastronarde, 2005), dose-symmetric tilt-series were acquired (Hagen et al., 2017), starting at +10° to match the pre-tilt of the lamella and then proceeding with 2° steps to roughly −50° and +70°. Individual tilts were recorded in movie mode with 10 frames per second, at an object pixel size of 3.52 Å and a target defocus of −4 to −5.5 μm. The total accumulated dose deposited on each tilt-series ranged from 90 to 140 e-/Å^2^. Each tomogram was acquired from a separate cell and thus is both a biological and technical replicate.

### *In situ* cryo-ET reconstruction, segmentation, and image analysis

The tilt-series were preprocessed and assembled with the STOPGAP tomoman pipeline (https://github.com/williamnwan/STOPGAP), including drift correction of raw frames with MotionCor2 software (Zheng et al., 2017) and dose-weighting of individual tilts. Using IMOD software (Kremer et al., 1996), the assembled tilt-series were aligned with patch tracking, and bin4 tomographic volumes (14.08 Å pixel size) were reconstructed by weighted back projection. To enhance contrast for display, we applied the tom_deconv deconvolution filter (https://github.com/dtegunov/tom_deconv) to the bin4 tomograms. Slices through tomogram volumes (Figs. 4–6, S14) were generated with the IMOD 3dmod viewer. Tomogram segmentation was performed in Amira software (FEI, Thermo Fisher Scientific), using a combination of manual segmentation and automated membrane detection from the TomoSegMemTV software package (Martinez-Sanchez et al., 2014).

Using the IMOD 3dmod viewer, 2D slices (zoomed 6x with interpolation) were generated from the bin4 tomographic volumes. The slices were oriented with the xy-plane perpendicular to the thylakoid membranes. Using Fiji software (Schindelin et al., 2012), line scan intensity profiles were generated with a 40 pixel-wide line drawn perpendicular to the thylakoid sheet. Lumen width was measured between the shoulders of the peaks corresponding to opposing lipid bilayers of each thylakoid (see Engel et al. 2015). The measurement point on the peak shoulder was defined as the halfway point between the dark peak of the EM density and the adjacent light peak of the fridge caused by the contrast transfer function. Similarly, membrane thickness was estimated by measuring the width of the peak corresponding to each bilayer (from shoulder to shoulder). A total of 50 tomograms were reconstructed, from which 48 were used for the analysis (WT: 7 LL and 6 HL; F4E mutant: 12 LL and 9 HL; V11E mutant: 7 LL and 7 HL). A total of 283 thylakoid membrane regions were used to generate 346 line scans (WT: 37 scans on 26 LL thylakoids and 28 scans on 25 HL thylakoids; F4E: 87 scans on 71 LL thylakoids and 63 scans on 52 HL thylakoids; V11E: 63 scans on 51 LL thylakoids and 68 scans on 58 HL thylakoids). For each thylakoid, scans were performed at the widest region and not more than once per 500 nm of thylakoid length.

## Author Contributions

Cryo-EM data acquisition and structure determination were performed by T.K.G., with guidance from J.M.S. Molecular modeling was performed by S.K., with guidance from T.R. Cloning and production of VIPP1 proteins was performed by K.G. and J. Niemeyer, with guidance from M. Schroda. Creation and physiological characterization of the *Synechocystis* VIPP1 mutants was performed by S.H., with guidance from J. Nickelsen. *In situ* cryo-ET data acquisition and analysis was performed by W.W. and S.K., with help from M. Schaffer, A.R., and B.D.E. Additional protein purification and nucleotide hydrolysis assays were performed by N.O., with guidance from W.S. M. Strauss. aided with preliminary EM image analysis. W.B. and J.M.P. provided guidance, financial support and access to instrumentation. T.K.G., J.M.S., M. Schroda, and B.D.E. wrote the paper, with input from all authors.

## Data Availability

Cryo-EM maps will be made available in the Electron Microscopy Data Bank (EMDB). Atomic models will be made available in the Protein Data Bank (PDB).

## Acknowledgements

We thank Elisabeth Weyher-Stingl and Victoria Sanchez Caballero at the MPI Biochemistry core facility for assistance with mass spectrometry, Günter Pfeifer and Florian Beck for help with microscopy and image analysis, Gert Bange and Ricardo Righetto for helpful discussions, and Karin Engel for critically reading the manuscript. This work was supported by grants from the Deutsche Forschungsgemeinschaft to J.N. (NI 390/9-2 as part of FOR2092), J.M.S (Emmy Noether grant SCHU 3364/1-1), M.S. (Schr 617/8-2 as part of FOR2092, TRR 175 project C02), and B.D.E. (EN 1194/1-1 as part of FOR2092), as well as KAKENHI grants to W.S. (16H06554 from the Ministry of Education, Culture, Sports, Science and Technology, 17H03699 from the Japanese Society for the Promotion of Science). Additional funding was provided by the Max Planck Society, the Helmholtz Zentrum München, and by LMU Munich’s Institutional Strategy “LMU Excellent” within the framework of the German Excellence Initiative.

## References

Asano, S., Engel, B.D., and Baumeister, W. (2016). In situ cryo-electron tomography: A post-reductionist approach to structural biology. J Mol Biol. 428: 332–343.

Aseeva, E., Ossenbuhl, F., Eichacker, L.A., Wanner, G., Soll, J., and Vothknecht, U.C. (2004). Complex formation of vipp1 depends on its alpha-helical pspa-like domain. J Biol Chem. 279: 35535–35541.

Aseeva, E., Ossenbuhl, F., Sippel, C., Cho, W.K., Stein, B., Eichacker, L.A., Meurer, J., Wanner, G., Westhoff, P., Soll, J., and Vothknecht, U.C. (2007). Vipp1 is required for basic thylakoid membrane formation but not for the assembly of thylakoid protein complexes. Plant Physiol Biochem. 45: 119–128.

Banerjee, S., Bartesaghi, A., Merk, A., Rao, P., Bulfer, S.L., Yan, Y., Green, N., Mroczkowski, B., Neitz, R.J., Wipf, P., Falconieri, V., Deshaies, R.J., Milne, J.L., Huryn, D., Arkin, M., and Subramaniam, S. (2016). 2.3 a resolution cryo-em structure of human p97 and mechanism of allosteric inhibition. Science. 351: 871–875.

Bryan, S.J., Burroughs, N.J., Shevela, D., Yu, J., Rupprecht, E., Liu, L.-N., Mastroianni, G., Xue, Q., Llorente-Garcia, I., Leake, M.C., Eichacker, L.A., Schneider, D., Nixon, P.J., and Mullineaux, C.W. (2014). Localisation and interactions of the vipp1 protein in cyanobacteria. Molecular Microbiology. 94: 1179–1195.

Cardone, G., Heymann, J.B., and Steven, A.C. (2013). One number does not fit all: Mapping local variations in resolution in cryo-em reconstructions. Journal of structural biology. 184: 226–236.

Chauhan, J.S., Mishra, N.K., and Raghava, G.P. (2009). Identification of atp binding residues of a protein from its primary sequence. BMC Bioinformatics. 10: 434.

Chauhan, J.S., Mishra, N.K., and Raghava, G.P.S. (2010). Prediction of gtp interacting residues, dipeptides and tripeptides in a protein from its evolutionary information. BMC bioinformatics. 11: 301–301.

Darwin, A.J. (2013). Stress relief during host infection: The phage shock protein response supports bacterial virulence in various ways. PLoS pathogens. 9: e1003388–e1003388.

DeLisa, M.P., Lee, P., Palmer, T., and Georgiou, G. (2004). Phage shock protein pspa of <em>escherichia coli</em> relieves saturation of protein export via the tat pathway. Journal of Bacteriology. 186: 366–373.

Denais, C.M., Gilbert, R.M., Isermann, P., McGregor, A.L., te Lindert, M., Weigelin, B., Davidson, P.M., Friedl, P., Wolf, K., and Lammerding, J. (2016). Nuclear envelope rupture and repair during cancer cell migration. Science. 352: 353–358.

Eaton-Rye, J.J. (2004). The construction of gene knockouts in the cyanobacterium synechocystis sp. Pcc 6803. Methods Mol Biol. 274: 309–324.

Fuhrmann, E., Bultema, J.B., Kahmann, U., Rupprecht, E., Boekema, E.J., and Schneider, D. (2009a). The vesicle-inducing protein 1 from synechocystis sp. Pcc 6803 organizes into diverse higher-ordered ring structures. Mol Biol Cell. 20: 4620–4628.

Fuhrmann, E., Gathmann, S., Rupprecht, E., Golecki, J., and Schneider, D. (2009b). Thylakoid membrane reduction affects the photosystem stoichiometry in the cyanobacterium synechocystis sp. Pcc 6803. Plant Physiology. 149: 735–744.

Gat, Y., Schuller, J.M., Lingaraju, M., Weyher, E., Bonneau, F., Strauss, M., Murray, P.J., and Conti, E. (2019). Insp6 binding to pikk kinases revealed by the cryo-em structure of an smg1-smg8-smg9 complex. Nat Struct Mol Biol. 26: 1089–1093.

Goddard, T.D., Huang, C.C., Meng, E.C., Pettersen, E.F., Couch, G.S., Morris, J.H., and Ferrin, T.E. (2018). Ucsf chimerax: Meeting modern challenges in visualization and analysis. Protein science : a publication of the Protein Society. 27: 14–25.

Goh, B.C., Hadden, J.A., Bernardi, R.C., Singharoy, A., McGreevy, R., Rudack, T., Cassidy, C.K., and Schulten, K. (2016). Computational methodologies for real-space structural refinement of large macromolecular complexes. Annual Review of Biophysics. 45: 253–278.

Gutu, A., Chang, F., and O’Shea, E.K. (2018). Dynamical localization of a thylakoid membrane binding protein is required for acquisition of photosynthetic competency. Molecular microbiology. 108: 16–31.

Hagen, W.J.H., Wan, W., and Briggs, J.A.G. (2017). Implementation of a cryo-electron tomography tilt-scheme optimized for high resolution subtomogram averaging. J Struct Biol. 197: 191–198.

Hamelryck, T., and Manderick, B. (2003). Pdb file parser and structure class implemented in python. Bioinformatics. 19: 2308–2310.

Hankamer, B.D., Elderkin, S.L., Buck, M., and Nield, J. (2004). Organization of the aaa(+) adaptor protein pspa is an oligomeric ring. J Biol Chem. 279: 8862–8866.

Heinz, S., Rast, A., Shao, L., Gutu, A., Gugel, I.L., Heyno, E., Labs, M., Rengstl, B., Viola, S., Nowaczyk, M.M., Leister, D., and Nickelsen, J. (2016). Thylakoid membrane architecture in synechocystis depends on curt, a homolog of the granal curvature thylakoid1 proteins. Plant Cell.

Hennig, R., Heidrich, J., Saur, M., Schmuser, L., Roeters, S.J., Hellmann, N., Woutersen, S., Bonn, M., Weidner, T., Markl, J., and Schneider, D. (2015). Im30 triggers membrane fusion in cyanobacteria and chloroplasts. Nat Commun. 6: 7018.

Hennig, R., West, A., Debus, M., Saur, M., Markl, J., Sachs, J.N., and Schneider, D. (2017). The im30/vipp1 c-terminus associates with the lipid bilayer and modulates membrane fusion. Biochim Biophys Acta Bioenerg. 1858: 126–136.

Hirama, T., Lu, S.M., Kay, J.G., Maekawa, M., Kozlov, M.M., Grinstein, S., and Fairn, G.D. (2017). Membrane curvature induced by proximity of anionic phospholipids can initiate endocytosis. Nature Communications. 8: 1393.

Humphrey, W., Dalke, A., and Schulten, K. (1996). Vmd: Visual molecular dynamics. J Mol Graph. 14: 33–38, 27-38.

Hurley, J.H. (2015). Escrts are everywhere. The EMBO Journal. 34: 2398–2407.

Jimenez, A.J., Maiuri, P., Lafaurie-Janvore, J., Divoux, S., Piel, M., and Perez, F. (2014). Escrt machinery is required for plasma membrane repair. Science. 343: 1247136.

Jovanovic, G., Mehta, P., McDonald, C., Davidson, A.C., Uzdavinys, P., Ying, L., and Buck, M. (2014). The n-terminal amphipathic helices determine regulatory and effector functions of phage shock protein a (pspa) in escherichia coli. J Mol Biol. 426: 1498–1511.

Junglas, B., Siebenaller, C., Schlösser, L., Hellmann, N., and Schneider, D. (2020). Gtp hydrolysis by synechocystis im30 does not decisively affect its membrane remodeling activity. Scientific Reports. 10: 9793.

Kirchhoff, H., Hall, C., Wood, M., Herbstova, M., Tsabari, O., Nevo, R., Charuvi, D., Shimoni, E., and Reich, Z. (2011). Dynamic control of protein diffusion within the granal thylakoid lumen. Proc Natl Acad Sci U S A. 108: 20248–20253.

Kleerebezem, M., Crielaard, W., and Tommassen, J. (1996). Involvement of stress protein pspa (phage shock protein a) of escherichia coli in maintenance of the protonmotive force under stress conditions. The EMBO Journal. 15: 162–171.

Klotz, A., Georg, J., Bučinská, L., Watanabe, S., Reimann, V., Januszewski, W., Sobotka, R., Jendrossek, D., Hess, Wolfgang R., and Forchhammer, K. (2016). Awakening of a dormant cyanobacterium from nitrogen chlorosis reveals a genetically determined program. Current Biology. 26: 2862–2872.

Kobayashi, R., Suzuki, T., and Yoshida, M. (2007). Escherichia coli phage-shock protein a (pspa) binds to membrane phospholipids and repairs proton leakage of the damaged membranes. Molecular Microbiology. 66: 100–109.

Kremer, J.R., Mastronarde, D.N., and McIntosh, J.R. (1996). Computer visualization of three-dimensional image data using imod. J Struct Biol. 116: 71–76.

Kroll, D., Meierhoff, K., Bechtold, N., Kinoshita, M., Westphal, S., Vothknecht, U.C., Soll, J., and Westhoff, P. (2001). Vipp1, a nuclear gene of arabidopsis thaliana essential for thylakoid membrane formation. Proc Natl Acad Sci U S A. 98: 4238–4242.

Kunz, H.-H., Gierth, M., Herdean, A., Satoh-Cruz, M., Kramer, D.M., Spetea, C., and Schroeder, J.I. (2014). Plastidial transporters kea1, −2, and −3 are essential for chloroplast osmoregulation, integrity, and ph regulation in arabidopsis. Proceedings of the National Academy of Sciences. 111: 7480–7485.

Leaver-Fay, A., Tyka, M., Lewis, S.M., Lange, O.F., Thompson, J., Jacak, R., Kaufman, K.W., Renfrew, P.D., Smith, C.A., Sheffler, W., Davis, I.W., Cooper, S., Treuille, A., Mandell, D.J., Richter, F., Ban, Y.- E.A., Fleishman, S.J., Corn, J.E., Kim, D.E., Lyskov, S., Berrondo, M., Mentzer, S., Popović, Z., Havranek, J.J., Karanicolas, J., Das, R., Meiler, J., Kortemme, T., Gray, J.J., Kuhlman, B., Baker, D., and Bradley, P. (2011). Chapter nineteen - rosetta3: An object-oriented software suite for the simulation and design of macromolecules. In Methods in enzymology, M.L. Johnson, and L. Brand, eds. (Academic Press), pp. 545–574.

Li, H.-m., Kaneko, Y., and Keegstra, K. (1994). Molecular cloning of a chloroplastic protein associated with both the envelope and thylakoid membranes. Plant Molecular Biology. 25: 619–632.

Liebschner, D., Afonine, P.V., Baker, M.L., Bunkóczi, G., Chen, V.B., Croll, T.I., Hintze, B., Hung, L.W., Jain, S., McCoy, A.J., Moriarty, N.W., Oeffner, R.D., Poon, B.K., Prisant, M.G., Read, R.J., Richardson, J.S., Richardson, D.C., Sammito, M.D., Sobolev, O.V., Stockwell, D.H., Terwilliger, T.C., Urzhumtsev, A.G., Videau, L.L., Williams, C.J., and Adams, P.D. (2019). Macromolecular structure determination using x-rays, neutrons and electrons: Recent developments in phenix. Acta Crystallogr D Struct Biol. 75: 861–877.

Lindås, A.C., Karlsson, E.A., Lindgren, M.T., Ettema, T.J., and Bernander, R. (2008). A unique cell division machinery in the archaea. Proc Natl Acad Sci U S A. 105: 18942–18946.

Liu, C., Willmund, F., Golecki, J.R., Cacace, S., Hess, B., Markert, C., and Schroda, M. (2007). The chloroplast hsp70b-cdj2-cge1 chaperones catalyse assembly and disassembly of vipp1 oligomers in chlamydomonas. Plant J. 50: 265–277.

Manganelli, R., and Gennaro, M.L. (2017). Protecting from envelope stress: Variations on the phage-shock-protein theme. Trends in microbiology. 25: 205–216.

Martinez-Sanchez, A., Garcia, I., Asano, S., Lucic, V., and Fernandez, J.J. (2014). Robust membrane detection based on tensor voting for electron tomography. J Struct Biol. 186: 49–61.

Mastronarde, D.N. (2005). Automated electron microscope tomography using robust prediction of specimen movements. J Struct Biol. 152: 36–51.

McCullough, J., Clippinger, A.K., Talledge, N., Skowyra, M.L., Saunders, M.G., Naismith, T.V., Colf, L.A., Afonine, P., Arthur, C., Sundquist, W.I., Hanson, P.I., and Frost, A. (2015). Structure and membrane remodeling activity of escrt-iii helical polymers. Science (New York, NY). 350: 1548–1551.

McDonald, C., Jovanovic, G., Ces, O., and Buck, M. (2015). Membrane stored curvature elastic stress modulates recruitment of maintenance proteins pspa and vipp1. mBio. 6: e01188–01115.

McDonald, C., Jovanovic, G., Wallace, B.A., Ces, O., and Buck, M. (2017). Structure and function of pspa and vipp1 n-terminal peptides: Insights into the membrane stress sensing and mitigation. Biochimica et Biophysica Acta (BBA) - Biomembranes. 1859: 28–39.

Michaud-Agrawal, N., Denning, E.J., Woolf, T.B., and Beckstein, O. (2011). Mdanalysis: A toolkit for the analysis of molecular dynamics simulations. J Comput Chem. 32: 2319–2327.

Morris, G.M., Huey, R., Lindstrom, W., Sanner, M.F., Belew, R.K., Goodsell, D.S., and Olson, A.J. (2009). Autodock4 and autodocktools4: Automated docking with selective receptor flexibility. Journal of computational chemistry. 30: 2785–2791.

Murakami, S., and Packer, L. (1970). Protonation and chloroplast membrane structure. The Journal of cell biology. 47: 332–351.

Nguyen, H.C., Talledge, N., McCullough, J., Sharma, A., Moss, F.R., 3rd, Iwasa, J.H., Vershinin, M.D., Sundquist, W.I., and Frost, A. (2020). Membrane constriction and thinning by sequential escrt-iii polymerization. Nat Struct Mol Biol. 27: 392–399.

Nordhues, A., Schottler, M.A., Unger, A.K., Geimer, S., Schonfelder, S., Schmollinger, S., Rutgers, M., Finazzi, G., Soppa, B., Sommer, F., Muhlhaus, T., Roach, T., Krieger-Liszkay, A., Lokstein, H., Crespo, J.L., and Schroda, M. (2012). Evidence for a role of vipp1 in the structural organization of the photosynthetic apparatus in chlamydomonas. Plant Cell. 24: 637–659.

Ohnishi, N., Zhang, L., and Sakamoto, W. (2018). Vipp1 involved in chloroplast membrane integrity has gtpase activity in vitro. Plant Physiology. 177: 328–338.

Olmos, Y., Hodgson, L., Mantell, J., Verkade, P., and Carlton, J.G. (2015). Escrt-iii controls nuclear envelope reformation. Nature. 522: 236–239.

Osadnik, H., Schopfel, M., Heidrich, E., Mehner, D., Lilie, H., Parthier, C., Risselada, H.J., Grubmuller, H., Stubbs, M.T., and Bruser, T. (2015). Pspf-binding domain pspa1-144 and the pspa.F complex: New insights into the coiled-coil-dependent regulation of aaa+ proteins. Mol Microbiol. 98: 743–759.

Otters, S., Braun, P., Hubner, J., Wanner, G., Vothknecht, U.C., and Chigri, F. (2013). The first alpha-helical domain of the vesicle-inducing protein in plastids 1 promotes oligomerization and lipid binding. Planta. 237: 529–540.

Pettersen, E.F., Goddard, T.D., Huang, C.C., Couch, G.S., Greenblatt, D.M., Meng, E.C., and Ferrin, T.E. (2004). Ucsf chimera--a visualization system for exploratory research and analysis. J Comput Chem. 25: 1605–1612.

Phillips, J.C., Braun, R., Wang, W., Gumbart, J., Tajkhorshid, E., Villa, E., Chipot, C., Skeel, R.D., Kalé, L., and Schulten, K. (2005). Scalable molecular dynamics with namd. Journal of Computational Chemistry. 26: 1781–1802.

Punjani, A., Rubinstein, J.L., Fleet, D.J., and Brubaker, M.A. (2017). Cryosparc: Algorithms for rapid unsupervised cryo-em structure determination. Nat Methods. 14: 290–296.

Raab, M., Gentili, M., de Belly, H., Thiam, H.R., Vargas, P., Jimenez, A.J., Lautenschlaeger, F., Voituriez, R., Lennon-Duménil, A.M., Manel, N., and Piel, M. (2016). Escrt iii repairs nuclear envelope ruptures during cell migration to limit DNA damage and cell death. Science. 352: 359–362.

Rast, A., Heinz, S., and Nickelsen, J. (2015). Biogenesis of thylakoid membranes. Biochim Biophys Acta.

Rast, A., Schaffer, M., Albert, S., Wan, W., Pfeffer, S., Beck, F., Plitzko, J.M., Nickelsen, J., and Engel, B.D. (2019). Biogenic regions of cyanobacterial thylakoids form contact sites with the plasma membrane. Nat Plants. 5: 436–446.

Rippka, R., Deruelles, J., Waterbury, J.B., Herdman, M., and Stanier, R.Y. (1979). Generic assignments, strain histories and properties of pure cultures of cyanobacteria. Microbiology. 111: 1–61.

Rosenthal, P.B., and Henderson, R. (2003). Optimal determination of particle orientation, absolute hand, and contrast loss in single-particle electron cryomicroscopy. J Mol Biol. 333: 721–745.

Sali, A., and Blundell, T.L. (1993). Comparative protein modelling by satisfaction of spatial restraints. J Mol Biol. 234: 779–815.

Samson, R.Y., Obita, T., Freund, S.M., Williams, R.L., and Bell, S.D. (2008). A role for the escrt system in cell division in archaea. Science. 322: 1710–1713.

Saraste, M., Sibbald, P.R., and Wittinghofer, A. (1990). The p-loop--a common motif in atp- and gtp-binding proteins. Trends Biochem Sci. 15: 430–434.

Saur, M., Hennig, R., Young, P., Rusitzka, K., Hellmann, N., Heidrich, J., Morgner, N., Markl, J., and Schneider, D. (2017). A janus-faced im30 ring involved in thylakoid membrane fusion is assembled from im30 tetramers. Structure. 25: 1380–1390.e1385.

Schaffer, M., Engel, B.D., Laugks, T., Mahamid, J., Plitzko, J.M., and Baumeister, W. (2015). Cryo-focused ion beam sample preparation for imaging vitreous cells by cryo-electron tomography. Bio Protoc. 5: e1575.

Schaffer, M., Mahamid, J., Engel, B.D., Laugks, T., Baumeister, W., and Plitzko, J.M. (2017). Optimized cryo-focused ion beam sample preparation aimed at in situ structural studies of membrane proteins. J Struct Biol. 197: 73–82.

Scheffer, L.L., Sreetama, S.C., Sharma, N., Medikayala, S., Brown, K.J., Defour, A., and Jaiswal, J.K. (2014). Mechanism of ca²⁺-triggered escrt assembly and regulation of cell membrane repair. Nat Commun. 5: 5646.

Scheres, S.H.W. (2012). Relion: Implementation of a bayesian approach to cryo-em structure determination. Journal of Structural Biology. 180: 519–530.

Schindelin, J., Arganda-Carreras, I., Frise, E., Kaynig, V., Longair, M., Pietzsch, T., Preibisch, S., Rueden, C., Saalfeld, S., Schmid, B., Tinevez, J.Y., White, D.J., Hartenstein, V., Eliceiri, K., Tomancak, P., and Cardona, A. (2012). Fiji: An open-source platform for biological-image analysis. Nat Methods. 9: 676–682.

Schottkowski, M., Gkalympoudis, S., Tzekova, N., Stelljes, C., Schunemann, D., Ankele, E., and Nickelsen, J. (2009). Interaction of the periplasmic prata factor and the psba (d1) protein during biogenesis of photosystem ii in synechocystis sp. Pcc 6803. J Biol Chem. 284: 1813–1819.

Schweitzer, A., Aufderheide, A., Rudack, T., Beck, F., Pfeifer, G., Plitzko, J.M., Sakata, E., Schulten, K., Förster, F., and Baumeister, W. (2016). Structure of the human 26s proteasome at a resolution of 3.9 å. Proceedings of the National Academy of Sciences. 113: 7816–7821.

Siebenaller, C., Junglas, B., Lehmann, A., Hellmann, N., and Schneider, D. (2020). Proton leakage is sensed by im30 and activates im30-triggered membrane fusion. Int J Mol Sci. 21.

Spetea, C., and Schoefs, B. (2010). Solute transporters in plant thylakoid membranes: Key players during photosynthesis and light stress. Commun Integr Biol. 3: 122–129.

Staehelin, L.A., Giddings, T.H., Jr., Kiss, J.Z., and Sack, F.D. (1990). Macromolecular differentiation of golgi stacks in root tips of arabidopsis and nicotiana seedlings as visualized in high pressure frozen and freeze-substituted samples. Protoplasma. 157: 75–91.

Stengel, A., Gugel, I.L., Hilger, D., Rengstl, B., Jung, H., and Nickelsen, J. (2012). Initial steps of photosystem ii de novo assembly and preloading with manganese take place in biogenesis centers in synechocystis. Plant Cell. 24: 660–675.

Theis, J., Gupta, T.K., Klingler, J., Wan, W., Albert, S., Keller, S., Engel, B.D., and Schroda, M. (2019). Vipp1 rods engulf membranes containing phosphatidylinositol phosphates. Sci Rep. 9: 8725.

Trabuco, L.G., Villa, E., Mitra, K., Frank, J., and Schulten, K. (2008). Flexible fitting of atomic structures into electron microscopy maps using molecular dynamics. Structure. 16: 673–683.

Tsunekawa, K., Shijuku, T., Hayashimoto, M., Kojima, Y., Onai, K., Morishita, M., Ishiura, M., Kuroda, T., Nakamura, T., Kobayashi, H., Sato, M., Toyooka, K., Matsuoka, K., Omata, T., and Uozumi, N. (2009). Identification and characterization of the na+/h+ antiporter nhas3 from the thylakoid membrane of synechocystis sp. Pcc 6803. J Biol Chem. 284: 16513–16521.

van der Does, C., Swaving, J., van Klompenburg, W., and Driessen, A.J.M. (2000). Non-bilayer lipids stimulate the activity of the reconstituted bacterial protein translocase. Journal of Biological Chemistry. 275: 2472–2478.

Vietri, M., Schink, K.O., Campsteijn, C., Wegner, C.S., Schultz, S.W., Christ, L., Thoresen, S.B., Brech, A., Raiborg, C., and Stenmark, H. (2015). Spastin and escrt-iii coordinate mitotic spindle disassembly and nuclear envelope sealing. Nature. 522: 231–235.

Vothknecht, U.C., Otters, S., Hennig, R., and Schneider, D. (2011). Vipp1: A very important protein in plastids?! Journal of Experimental Botany.

Wagner, T., Merino, F., Stabrin, M., Moriya, T., Antoni, C., Apelbaum, A., Hagel, P., Sitsel, O., Raisch, T., Prumbaum, D., Quentin, D., Roderer, D., Tacke, S., Siebolds, B., Schubert, E., Shaikh, T.R., Lill, P., Gatsogiannis, C., and Raunser, S. (2019). Sphire-cryolo is a fast and accurate fully automated particle picker for cryo-em. Commun Biol. 2: 218.

Walker, J.E., Saraste, M., Runswick, M.J., and Gay, N.J. (1982). Distantly related sequences in the alpha- and beta-subunits of atp synthase, myosin, kinases and other atp-requiring enzymes and a common nucleotide binding fold. Embo j. 1: 945–951.

Wehmer, M., Rudack, T., Beck, F., Aufderheide, A., Pfeifer, G., Plitzko, J.M., Forster, F., Schulten, K., Baumeister, W., and Sakata, E. (2017). Structural insights into the functional cycle of the atpase module of the 26s proteasome. Proc Natl Acad Sci U S A. 114: 1305–1310.

Westphal, S., Heins, L., Soll, J., and Vothknecht, U.C. (2001). Vipp1 deletion mutant of synechocystis: A connection between bacterial phage shock and thylakoid biogenesis? Proc Natl Acad Sci U S A. 98: 4243–4248.

Willmund, F., Mühlhaus, T., Wojciechowska, M., and Schroda, M. (2007). The nh2-terminal domain of the chloroplast grpe homolog cge1 is required for dimerization and cochaperone function in vivo. J Biol Chem. 282: 11317–11328.

Zhang, K. (2016). Gctf: Real-time ctf determination and correction. J Struct Biol. 193: 1–12.

Zhang, L., Kato, Y., Otters, S., Vothknecht, U.C., and Sakamoto, W. (2012). Essential role of vipp1 in chloroplast envelope maintenance in arabidopsis. The Plant Cell. 24: 3695–3707.

Zhang, L., Kondo, H., Kamikubo, H., Kataoka, M., and Sakamoto, W. (2016). Vipp1 has a disordered c-terminal tail necessary for protecting photosynthetic membranes against stress. Plant Physiology. 171: 1983–1995.

Zhang, S., Shen, G., Li, Z., Golbeck, J.H., and Bryant, D.A. (2014). Vipp1 is essential for the biogenesis of photosystem i but not thylakoid membranes in synechococcus sp. Pcc 7002.Journal of Biological Chemistry.

Zheng, S.Q., Palovcak, E., Armache, J.-P., Verba, K.A., Cheng, Y., and Agard, D.A. (2017). Motioncor2: Anisotropic correction of beam-induced motion for improved cryo-electron microscopy. Nat Meth. 14: 331–332.

Zivanov, J., Nakane, T., Forsberg, B.O., Kimanius, D., Hagen, W.J., Lindahl, E., and Scheres, S.H. (2018). New tools for automated high-resolution cryo-em structure determination in relion-3. Elife. 7.

Zivanov, J., Nakane, T., and Scheres, S.H.W. (2019). A bayesian approach to beam-induced motion correction in cryo-em single-particle analysis. IUCrJ. 6: 5–17.

